# Crosstalk between the glucocorticoid and mineralocorticoid receptor boosts glucocorticoid-induced killing of multiple myeloma cells

**DOI:** 10.1101/2023.05.10.540157

**Authors:** Dorien Clarisse, Stefan Prekovic, Philip Vlummens, Eleni Staessens, Karlien Van Wesemael, Jonathan Thommis, Daria Fijalkowska, Guillaume Acke, Wilbert Zwart, Ilse M. Beck, Fritz Offner, Karolien De Bosscher

**Affiliations:** VIB Center for Medical Biotechnology, Ghent, Belgium; Department of Biomolecular Medicine, Ghent University, Ghent, Belgium; Cancer Research Institute Ghent (CRIG), Ghent, Belgium; Center for Molecular Medicine, University Medical Center Utrecht, Utrecht, The Netherlands; Department of Internal Medicine and Pediatrics, Ghent University Hospital, Ghent, Belgium; Department of Chemistry, Ghent University, Ghent, Belgium; Division of Oncogenomics, Oncode Institute, The Netherlands Cancer Institute, Amsterdam, The Netherlands; Department of Health Sciences, Odisee University of Applied Sciences, Ghent, Belgium

## Abstract

The glucocorticoid receptor (GR) is a crucial drug target in multiple myeloma as its activation with glucocorticoids effectively triggers myeloma cell death. However, as high-dose glucocorticoids are also associated with deleterious side effects, novel approaches are urgently needed to improve GR action in myeloma. Here we reveal a functional crosstalk between GR and the mineralocorticoid receptor (MR) that culminates in improved myeloma cell killing. We show that the GR agonist Dexamethasone (Dex) downregulates MR levels in a GR-dependent way in myeloma cells. Co-treatment of Dex with the MR antagonist Spironolactone (Spi) enhances Dex-induced cell killing in primary, newly diagnosed GC-sensitive myeloma cells. In a relapsed GC-resistant setting, Spi alone induces distinct myeloma cell killing. On a mechanistic level, we find that a GR-MR crosstalk likely arises from an endogenous interaction between GR and MR in myeloma cells. Quantitative dimerization assays show that Spi reduces Dex-induced GR-MR heterodimerization and completely abolishes Dex-induced MR-MR homodimerization, while leaving GR-GR homodimerization intact. Unbiased transcriptomics analyses reveal that c-myc and many of its target genes are downregulated most by combined Dex-Spi treatment. Proteomics analyses further identify that several metabolic hallmarks are modulated most by this combination treatment. Finally, we identified a subset of Dex-Spi downregulated genes and proteins that may predict prognosis in the CoMMpass myeloma patient cohort. Our study demonstrates that GR-MR crosstalk is therapeutically relevant in myeloma as it provides novel strategies for glucocorticoid-based dose-reduction.

## Introduction

More than 10% of all patients with hematological malignancies are diagnosed with multiple myeloma, which is a plasma cell cancer that is localized in the bone marrow^1, 2^. Despite significant advances in myeloma treatment, synthetic glucocorticoids (GCs) such as Dexamethasone (Dex) remain an important pillar of the myeloma treatment protocol because of their strong anti-myeloma activities, justifying their continued use in all treatment stages^1, 3^. However, long-term use of high-dose GCs is hampered by the emergence of GC resistance and side effects including osteoporosis, hyperglycemia, muscle wasting and severe mood swings, which negatively impact patient quality-of-life and treatment adherence^4^.

Both the therapeutic and unwanted effects of GCs are exerted through ligand-mediated activation of the glucocorticoid receptor (GR); a transcription factor belonging to the superfamily of ligand-activated nuclear receptors^5^. Once GCs bind to GR, the receptor undergoes a conformational change that results in a rearrangement of the Hsp90-FKBP51-containing multi-protein complex that aids in nuclear translocation of GR^6, 7^. In the nucleus, ligand-activated GR can promote gene activation, which can be accomplished by GR homodimers binding to glucocorticoid response elements (GREs) of target gene promoters or enhancers^8, 9^. In contrast, GR monomers can trigger gene repression by interfering with gene expression programs of other DNA-bound transcription factors such as NF-κB and AP-1 via, for instance, a tethering mechanism^10^. However, the dominant interaction mode between GR oligomers and other transcription factors on DNA remains a topic of debate^11–15^.

The intricate interplay of GR oligomers with other nuclear receptor oligomers, also called nuclear receptor crosstalk, results in a unique gene expression profile that allows for a strengthening or weakening of each receptor’s activity^16^. This crosstalk was already established for GR and estrogen receptor α (ERα) in breast cancer and for GR and androgen receptor (AR) in prostate cancer^17^. We serendipitously found that besides GR, the structure-wise closely related mineralocorticoid receptor (MR) was differentially expressed between myeloma cell lines. The impact of a possible interplay between GR and MR on GC therapy responsiveness has however not been considered in myeloma.

MR responds to two physiological ligands, aldosterone and cortisol, in a cell-type dependent manner and is ubiquitously expressed^18, 19^. This receptor regulates the electrolyte balance and water homeostasis in epithelial cells, while in non-epithelial cells inappropriate MR activation triggers pro-inflammatory and profibrotic effects^20^. MR-mediated effects are counteracted by MR antagonists, such as Spironolactone (Spi), which are used in the clinic for their cardiovascular and renal protective functions and to lower blood pressure^21, 22^. Crosstalk mechanisms between GR and MR were shown in several tissues^16, 23, 24^ and result in the formation of GR-MR heterodimers or even higher order oligomers, thereby modulating the transcriptional activity of each receptor^23–27^. Several studies support that MR inhibits GR-mediated gene transcription following GR-MR heterodimerization^28–30^. However, a study in neuroblastoma cells shows that GC-induced transcription was enhanced by a tethering of MR to DNA-bound GR^23^. How the GC response in myeloma may be influenced by the interplay between GR and MR is still elusive.

In this study, we present a novel crosstalk mechanism between GR and MR in multiple myeloma cells that may offer a unique therapy-supportive angle for myeloma treatment. We show that GCs downregulate MR levels in a GR-dependent fashion and that inhibiting MR with Spi culminates in an enhanced Dex-induced myeloma cell killing. We further elaborate on this GR-MR crosstalk by showing that Spi reduces Dex-induced GR-MR heterodimerization and completely abolishes Dex-induced MR-MR homodimerization. Finally, we reveal the transcriptomic and proteomic signatures of the Dex-Spi combination treatment that underpin the enhanced myeloma cell killing effects and identify a subset of Dex-Spi-regulated targets that predict survival in the CoMMpass patient cohort.

## Results

### Dex downregulates MR levels in a GR-dependent way

We first examined whether GCs regulate MR mRNA and protein levels (Fig.1A,B) in five myeloma cell lines (MM1.S, OPM-2, L-363, U-266 and MM1.R)^31, 32^ with different sensitivities to Dex-mediated myeloma cell killing (Fig.1C). A 6h Dex treatment downregulated *NR3C2* (MR) transcripts in cells showing the highest GC-inducible MM cell killing (MM1.S, OPM-2 and L-363), while in cells with virtually no GC-mediated MM cell killing (U-266 and MM1.R), *NR3C2* mRNA levels remained unchanged (Fig.1A). MR protein levels were also downregulated in GC-sensitive MM1.S and OPM-2 cells following 24h Dex treatment and only slightly in GC-resistant GR-negative MM1.R cells (Fig.1B). Despite that *NR3C2* mRNA is present in all myeloma cell lines (Fig.1A), MR protein levels were hardly detectable in both U-266 and L-363 cells (Fig.1B).

**Fig. 1:**
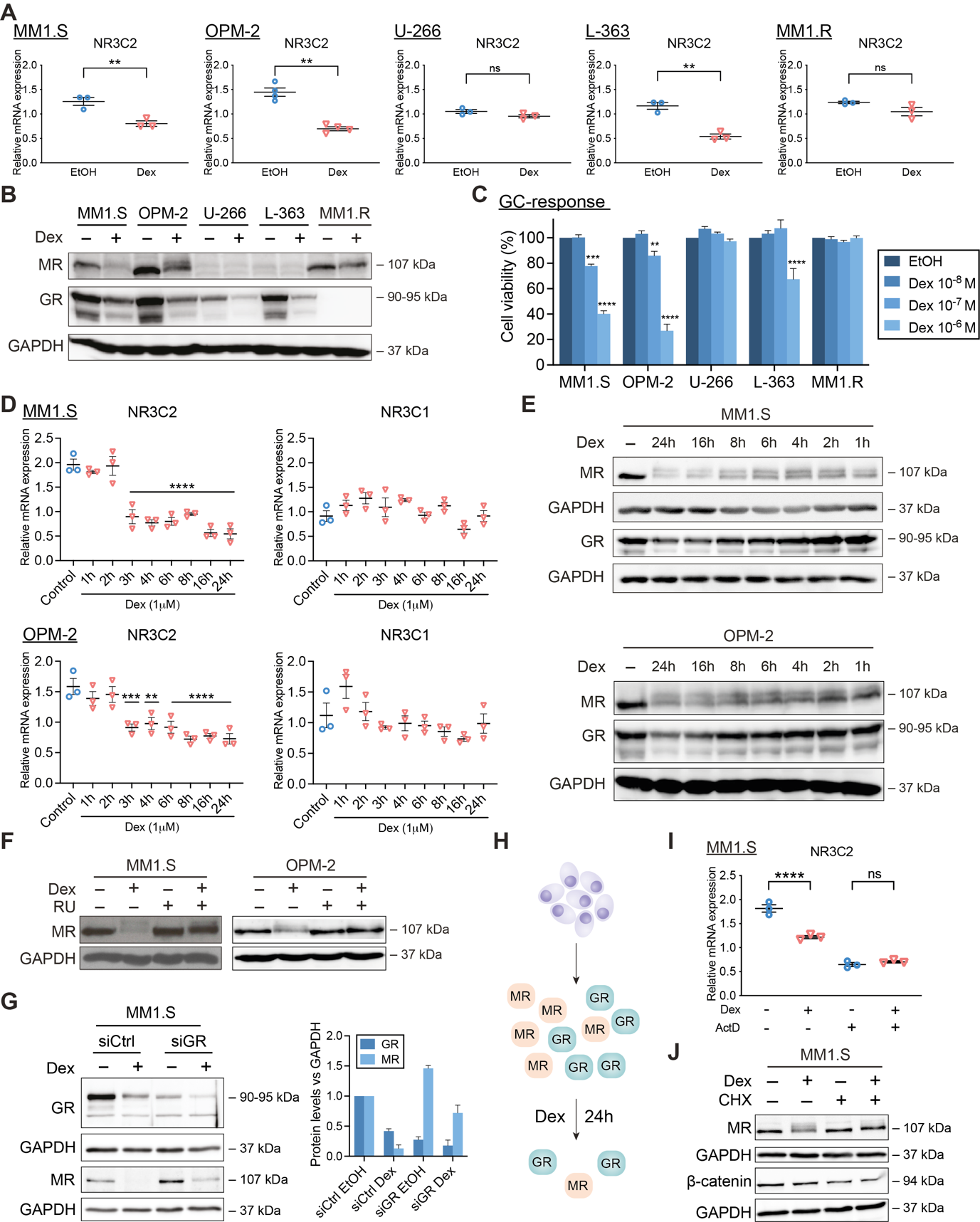
GCs downregulate MR mRNA and protein levels in a GR-dependent way. (**A-B**) MM1.S, OPM-2, U-266, L-363 and MM1.R cells were treated with Dex (10^−6^M) or solvent control (EtOH), (**A**) for 6h, followed by RT-qPCR (all N=3, except OPM-2: N=4), assessing the mRNA levels of *NR3C2*, or (**B**) for 24h, followed by WB analysis (N=3). The protein levels of MR (107kDa) and GR (90-95kDa) were determined, with GAPDH (37kDa) as loading control. (**C**) MM1.S, OPM-2, U-266, L-363 and MM1.R cells were treated for 72h with a Dex concentration range (10^−6^M-10^−8^M) or solvent control (EtOH, set as 100%), followed by a CelltiterGlo cell viability assay (72h Dex range recapitulated from Fig. 2I and 3B-D). The bar plots represent the mean +/− SEM. Statistical analyses were performed using GraphPad Prism 9, using a two-way ANOVA with post-hoc testing. Per cell line, 10^−6^M Dex and 10^−7^M Dex conditions were statistically compared to the 10^−8^M Dex condition. (**D-E**) MM1.S or OPM-2 cells were treated for different time points with Dex (10^−6^M) or solvent control (EtOH) followed by (**D**) RT-qPCR (N=3), assessing the mRNA levels of *NR3C2* and *NR3C1* and in which statistical analyses compared each time point to solvent control, or (**E**) WB analysis (N=3), in which the protein levels of MR (107kDa) and GR (90-95kDa) were determined, with GAPDH (37kDa) as loading control. (**F**) OPM-2 and MM1.S cells were treated with Dex (10^−6^M), RU (10^−5^M), a combination thereof or solvent control for 24h, followed by WB analysis (N=3). The protein levels of MR (107kDa) were determined, with GAPDH (37kDa) as loading control. (**G**) MM1.S cells were nucleofected with siCtrl (scrambled) or siGR and 48h post-nucleofection treated for another 24h with Dex (10^−6^M) or solvent control; followed by WB analysis (N=3) and band densitometric analysis (bar plot). The latter shows the normalized GR or MR protein levels (vs. GAPDH), averaged over 3 biological replicates. (**H**) Graphical summary. In MM cells containing GR and MR protein, Dex downregulates GR protein levels and to an even higher extent MR protein levels, especially at 24h. (**I**) MM1.S cells were treated for 3h with Dex (10^−6^M), ActD (1μg/mL), a combination thereof or solvent, followed by RT-qPCR (N=3), assessing the mRNA levels of *NR3C2*. (**J**) MM1.S cells were treated for 6h with Dex (10^−6^M), CHX (20μg/mL), a Dex/CHX combination or solvent control, followed by WB analysis (N=3) and band densitometric analysis. The protein levels of MR (107kDa), or β-catenin (94kDa; positive control for inhibition of protein translation) were determined, with GAPDH (37kDa) as loading control. Data information: (**A, D, I**) RT-qPCRs were analyzed using qBaseplus with *SDHA, RPL13A* and *YWHAZ* serving as reference genes. The scatter plots represent the mean (solid line) +/− SEM. Statistical analyses were performed using GraphPad Prism 9, using a one-way ANOVA with post-hoc testing. (**B, E, F, G, J**) One representative image is shown for each WB experiment, with the number of biological replicates mentioned in each panel description.

Dex decreased *NR3C1* (GR) mRNA levels only in L-363 cells and thus not in the GC-inducible MM1.S and OPM-2 cells (Supplementary Fig.S1; MM1.R is NR3C1-negative). GR protein, however, consistently underwent homologous downregulation following 24h Dex treatment (also known as negative feedback of GR) in all GR-containing MM cells (Fig.1B), which agrees with several reports^33–36^. Next to both receptors, we examined the Dex response of shared target genes. *TSC22D3* (GILZ)^37^ and *FKBP5*^31^ mRNA levels were upregulated by Dex in all MM cells except MM1.R cells, while *SGK1*^41^ mRNA levels are decreased by Dex in MM1.S, L-363 and U-266 cells (Supplementary Fig.S1).

The dynamic behavior of this Dex-induced MR downregulation was illustrated by showing that from 3h onwards, Dex significantly decreased *NR3C2* mRNA levels in MM1.S and OPM-2 cells; a fast regulation that was largely recapitulated at the protein level (Fig.1D-E). In contrast to *NR3C1* mRNA levels, Dex gradually declined GR protein levels over time (Fig.1D-E), as shown before^36^.

To confirm our observations across GCs with different potencies, we compared, ranked from high to low potency, the following ligands: Dex, fluocinolone acetonide (FA), prednisolone (Pred) and hydrocortisone (Hcort). All GCs consistently downregulated MR protein levels in MM1.S cells (Supplementary Fig.S2A). We observed a double MR band (with Dex, FA) and even multiple MR bands (with Pred, HCort) in MM1.S, while in MM1.R only Pred and HCort induced a clear double MR band, suggestive of post-translational modification of MR^38^.

Several lines of evidence support that the Dex-induced MR protein downregulation is largely GR-dependent. First, we used the GR antagonist RU486. A Dex/RU486 combination left MR protein levels intact compared to Dex alone in MM1.S and OPM-2 cells (Fig.1F). In addition, an siRNA-based GR knockdown in MM1.S cells showed that MR levels are at least partially protected from Dex-induced downregulation in siGR compared to siCtrl conditions (Fig.1G), overall supporting a GR-dependent mechanism. Noteworthy, knockdown of GR was already sufficient to increase the basal MR protein levels in MM1.S cells. Thirdly, in GR-negative MM1.R cells, Dex may bind MR instead, although this was clearly not sufficient to trigger the pronounced MR downregulation that was observed in GR-positive MM1.S cells.

To investigate whether Dex lowered *NR3C2* levels post-transcriptionally, we used actinomycin D (ActD) to block de novo transcription in MM1.S cells. ActD by itself reduced *NR3C2* levels three-fold compared to solvent condition, indicating that MR mRNA is unstable (Fig.1I). Addition of Dex on top of ActD could not further reduce the residual *NR3C2* mRNA levels, suggesting that novel gene transcription is needed. Next, we evaluated whether mechanisms centered at the protein level were contributing to Dex-induced MR downregulation. The protein translation inhibitor cycloheximide (CHX), of which the activity was confirmed via β-catenin downregulation (positive control), combined with Dex did not further reduce the MR protein levels at 6h treatment compared to Dex alone. This indicates that Dex requires novel protein synthesis to decrease MR protein levels (Fig.1J). In addition, Dex did not decline MR protein levels via lysosomal degradation, as assessed with chloroquine (positive control: LC-3) or via proteasomal degradation, as evaluated with the proteasome inhibitor MG132 (positive control: Hsp70) (Supplementary Fig.S2B-E).

Taken together, GCs decrease both MR mRNA and protein levels in MM cell lines with different degrees of GC-mediated MM cell killing in a GR-dependent manner, corroborating the existence of GR-MR crosstalk in these cells (Fig.1H). Our findings also demonstrate that MR mRNA is unstable and that Dex requires de novo transcription and translation to decrease MR levels in MM cells.

### MR antagonism enhances GC responsiveness of MM1.S cells

Because Dex can decrease MR mRNA and protein levels in MM cells, we examined whether a targeting of MR could affect the anti-MM activity of GCs. Hereto, we used three different strategies. First, we used MR knockdown using siRNA’s (siMR, Fig.2A-B) and found that the MM1.S cell viability was markedly reduced even in absence of Dex (Fig. 2A). Although the effect size by which Dex reduces the MM1.S cell viability was comparable in siCtrl and siMR conditions, the cell viability was significantly lower (∼55%) in the siMR Dex compared to the siCtrl Dex condition (∼80%, Fig.2A), Altogether, this suggests that MR presence may protect against myeloma cell death.

**Fig. 2:**
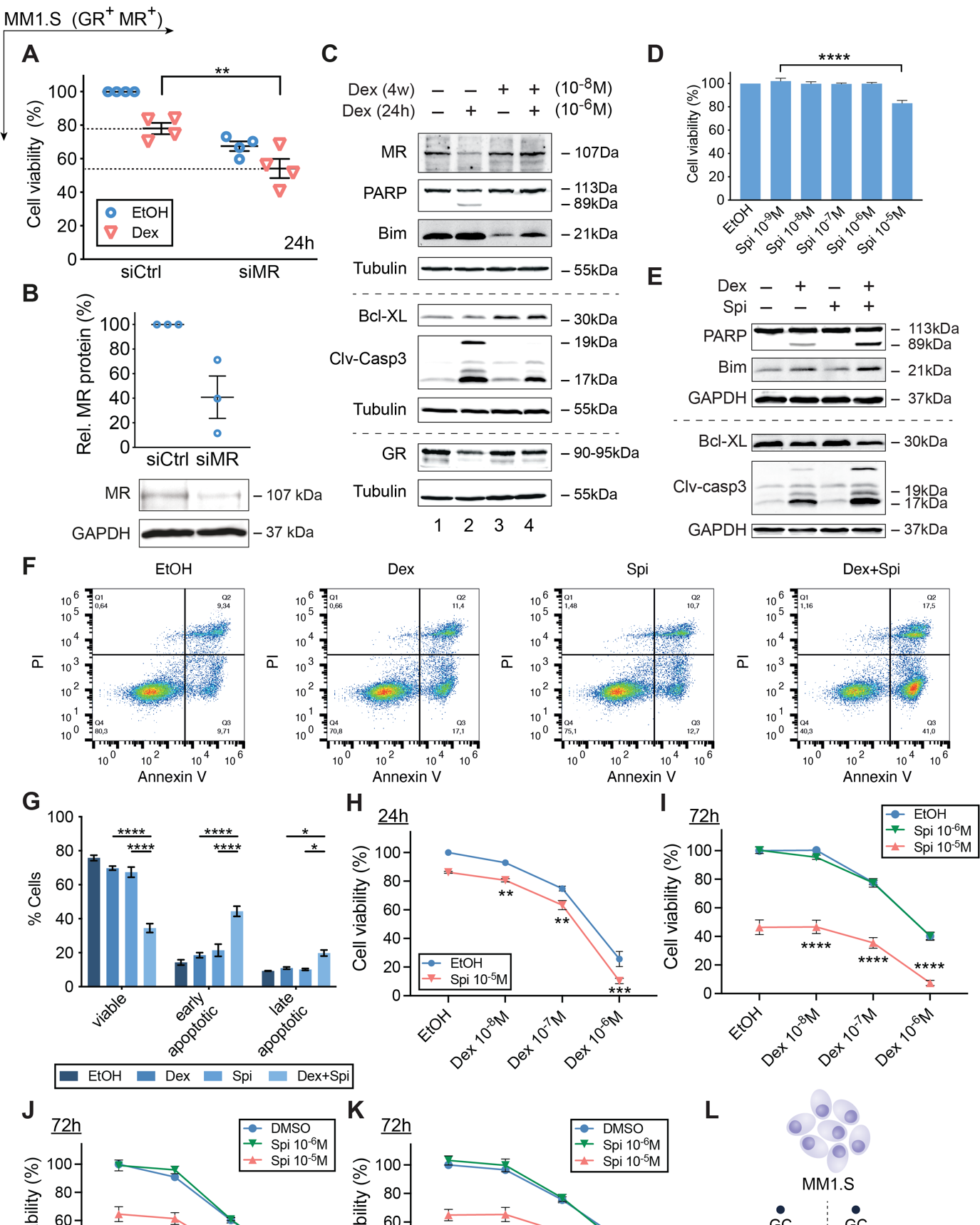
GC-induced MM1.S cell killing is promoted by the MR antagonist Spi. (**A**) MM1.S cells were nucleofected with siCtrl (scrambled) or siMR. 48h post-nucleofection, cells were reseeded and treated for another 24h with Dex (10^−6^M) or solvent control (EtOH), followed by a CelltiterGlo assay. The scatter plot represents the mean (solid line) +/− SEM (N=4). The siCtrl solvent condition was set as 100% and the other conditions were recalculated accordingly. (**B**) 72h post-nucleofection with siCtrl or siMR, WB analyses were performed and MR protein levels relative to GAPDH were quantified by band densitometric analysis using ImageJ. The scatter plot represents the mean +/− SEM (N=3). (**C**) MM1.S cells were treated for 4 weeks with 10^−8^M Dex (or EtOH), followed by 24h 10^−6^M Dex (or EtOH), and subjected to WB analyses (N=3). (**D**) MM1.S cells were treated with a Spi concentration range (10^−5^M-10^−9^M) or solvent control (set as 100%), followed by a CelltiterGlo assay (N=3). (**E-G**) MM1.S cells were treated with Dex (10^−6^M), Spi (10^−5^M) or a Dex-Spi combination for 24h, followed by (**E**) WB analyses (N=4) or (**F-G**) Annexin V/PI flow cytometric analyses (N=4). (**F**) Representative quadrant plots of 4 independent experiments for each treatment condition, with (**G**) bar plots showing the percentage of viable (Q4), early apoptotic (Q3), late apoptotic (Q2) averaged over all 4 biological repetitions +/− SEM. (**H-I**) MM1.S cells were treated with Dex (10^−6^M-10^−8^M), Spi (10^−5^-10^−6^M) or a Dex-Spi combination for (**H**) 24h (N=5) or (**I**) 72h (N=3), followed by a CelltiterGlo assay (solvent control set as 100%). (**J-K**) MM1.S cells were treated with Pred or Cort (10^−6^M-10^−8^M), Spi (10^−5^-10^−6^M) or a Pred/Spi or Cort/Spi combination for 72h (N=3), followed by a CelltiterGlo assay (solvent control set as 100%). (**L**) Summarizing model demonstrating that MR blockade increases Dex-induced MM1.S cell killing. Data information: (**A**, **D**, **F-J**) Statistical analyses were performed using GraphPad Prism 9, using (**A**, **D**) one-way or (**G-K**) two-way ANOVA with post-hoc testing. (**C, E**) Protein lysates were subjected to WB analyses, visualizing the protein levels of MR (107kDa), GR (90-95kDa), PARP (89 and 113kDa), Bim (21kDa), Bcl-XL (30kDa) and cleaved-caspase 3 (17 and 19 kDa). Tubulin (55kDa) or GAPDH (37kDa) served as loading controls. One representative image is shown for each WB experiment, with the number of biological replicates mentioned in each panel description.

Second, to evaluate how MR levels evolve upon prolonged GC treatment, we developed a cell model that mimics the gradual build-up of GC resistance. Here, MM1.S cells were treated for four weeks with a low dose of Dex (10^−8^M) followed by a high dose of Dex (10^−6^M) for 24h to assess the residual GC responsiveness of the MM cells to cell killing (Fig.2C). When MM cells were treated for four weeks with solvent, the additional 24h high-dose Dex resulted in a marked decrease in MR protein levels (Fig.2C, lane 1 vs 2), in line with Fig.1B. In contrast, after four weeks low-dose Dex, MR levels no longer declined following a 24h high-dose Dex (Fig.2C, lane 3 vs 4). Apoptotic marker analyses confirmed that the four weeks low-dose Dex, indeed rendered MM1.S cells refractory to the 24h high-dose Dex boost (Fig.2C, lane 2 vs 4). We found decreased cleavage of pro-apoptotic PARP and caspase 3, reduced levels of pro-apoptotic Bim and increased levels of anti-apoptotic Bcl-XL (Fig.2C, lane 2 vs 4).

Third, because MR knockdown promoted MM1.S cell killing (Fig.2A), we sought to complement these findings by using the MR antagonist Spironolactone (Spi). A concentration-response experiment showed that 10^−5^M Spi (24h) supported a mild cell killing in MM1.S cells (Fig.2D), but not in endothelial EA.hy926 cells (72h, toxicity control, Supplementary Fig.S3A). Next, we treated MM1.S cells for 24h with a combination of Dex (10^−6^M) and Spi (10^−5^M) and showed increased cleavage of pro-apoptotic PARP and caspase 3, and a decrease in anti-apoptotic Bcl-xL (Fig.2E). Confirmatory Annexin V/PI flow cytometric analyses showed that Dex-Spi combination decreased the percentage of viable MM1.S cells and increased the percentage of early-and late-apoptotic cells compared to each treatment alone (Fig.2F-G, Supplementary Fig.S3B). Spi enhanced Dex-induced MM1.S cell killing already at 24h (Fig.2H) and at 72h to an even higher extent (Fig.2I). Finally, also Pred- and Hcort-mediated MM1.S cell killing was boosted by Spi (Fig.2J-K).

Summarized, our results suggest that MR is a pro-survival factor in myeloma and that its pharmacological inhibition enhances GC-induced MM1.S cell killing.

### The ability of Spi to promote Dex-induced cell killing correlates with the Dex responsiveness of MM cell lines

To determine whether a Dex-Spi combination is effective across MM cell line models, we screened four other MM cell lines in which GCs induce MM cell killing to varying extents (Fig.1C). Whereas the MR antagonist Spi did not readily promote GC-mediated OPM-2 killing at 24h of treatment, this was observed after 72h of treatment (Fig.3A) as strongly as observed for MM1.S cells (Fig.2I). Hence, the threshold to obtain an efficient Dex-Spi-induced killing could have a time-dependent component when comparing OPM-2 to MM1.S.

Next, we tested MM cells that are less (or un)responsive to Dex in terms of cell killing to evaluate whether Dex-Spi still offers therapeutic benefit. Although L-363 cells respond slightly to Dex treatment, Spi did not significantly impact Dex-mediated cell killing of these cells after 72h (Fig.3B), which may be due to the very low amounts of MR that these cells contain (only detectable at mRNA level, Fig.1A-B). The same reasoning applies to the Dex-unresponsive U-266 cells, where Spi alone caused only a mild drop in cell viability (∼10%, Fig.3C). Interestingly, in MM1.R cells, which are GR-negative yet strongly MR-positive (Fig.1B), Spi alone triggered a marked decrease in cell viability at 72h of treatment (drop of ∼30%, Fig.3D), for which GR presence is clearly not required.

**Fig. 3:**
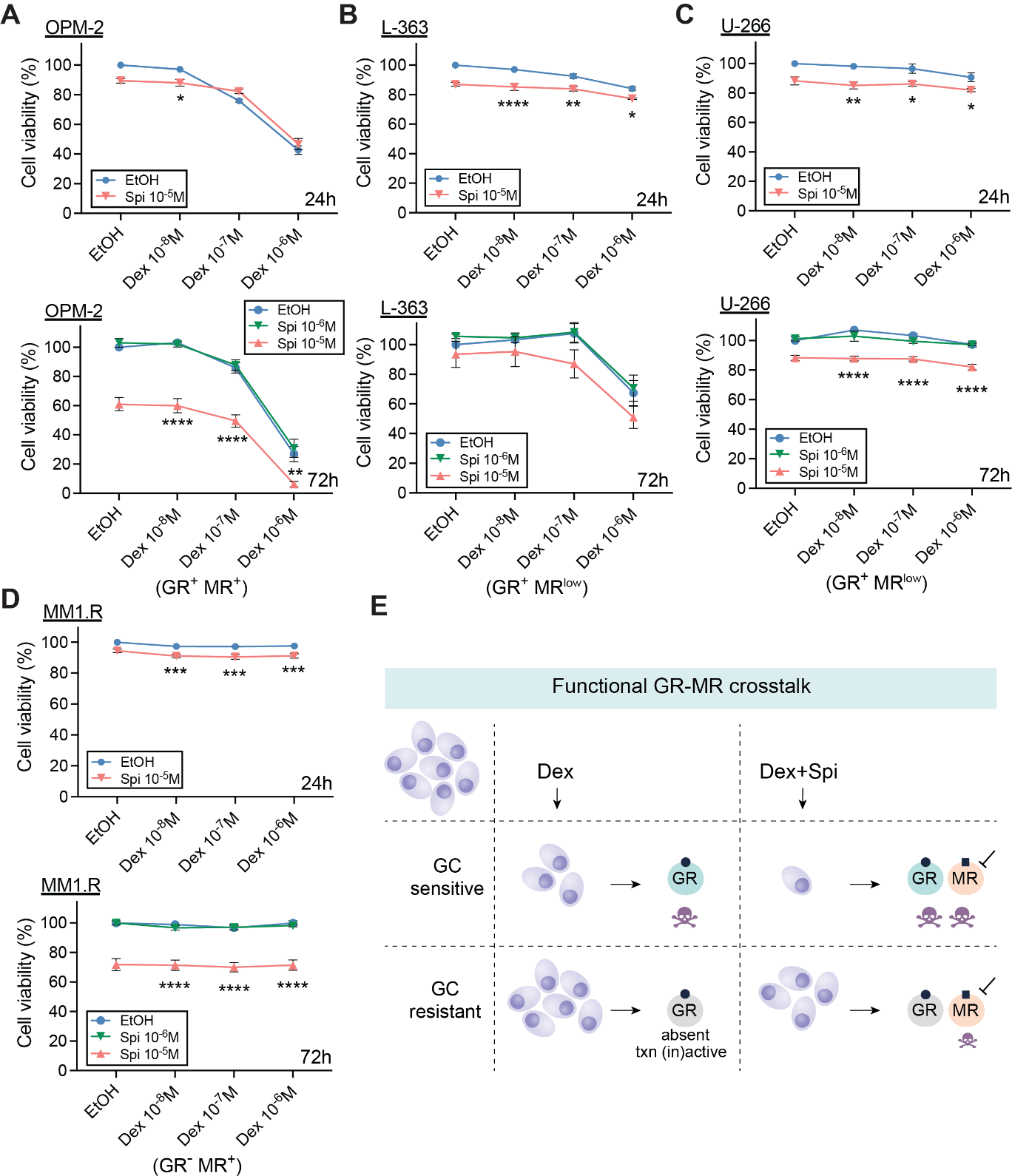
The MR antagonist Spi promotes cell killing of MM cells with varying degrees of Dex responsiveness. (**A-D**) Different myeloma cell lines including (**A**) OPM-2, (**B**) L-363, (**C**) U-266 and (**D**) MM1.R cells were treated with Dex (10^−6^M-10^−8^M), Spi (10^−5^-10^−6^M), a Dex-Spi combination or solvent control (set as 100%) for 24h or 72h, followed by a CelltiterGlo assay. Biological replicates: OPM-2 (24h N=6; 72h N=4), L-363 (24h and 72h N=3), U-266 (24h N=4, 72h N=3) and MM1.R (24h N=4, 72h N=3). (**E**) Graphical summary highlighting the existence of a functional crosstalk between GR and MR in MM cells. In GC-sensitive MM cells containing GR, Dex induces MM cell killing, which is further enhanced by the addition of Spi. In GC-resistant cells, where GR is either absent or transcriptionally (in)active, Dex loses its anti-MM activity, while Spi addition does trigger significant MM cell killing. Data information: (**A**-**D**) Statistical analyses were performed using GraphPad Prism 9 using two-way ANOVA with post-hoc testing.

In summary, in myeloma cells that contain detectable protein levels of both GR and MR, Spi enhances Dex-induced myeloma cell killing (Fig.3E).

### Combining lower doses of GC with MR antagonist enhances cell death of primary MM cells

To validate the potential of a Dex-Spi combination treatment in a preclinical context, we isolated primary MM cells from bone marrow aspirates of 10 MM patients at different disease stages (Table 1). In line with our observations in MM1.S and OPM-2 cells, newly diagnosed MM1 (Fig.4A) and MM2 (Fig.4B) patient cells as well as the premalignant smoldering MM9 patient cells (Fig.4K) displayed higher cell killing when combining Dex and Spi versus Dex alone. Importantly, 10^−7^M Dex combined with 10^−5^M Spi was at least equally efficacious as 10^−6^M Dex (∼40mg comparator dose in patients) alone (full arrow). As could be expected from a heterogeneous disease as myeloma, not all patient samples responded alike. Newly diagnosed MM3 patient cells (Fig.4C) strongly responded to Dex but had no additional benefit from Spi treatment. Furthermore, MM10 cells of a premalignant MGUS patient (Fig.4L) hardly responded to Dex treatment, while Spi alone reduced the cell viability with about 20% (dashed arrow). Notable, in all relapsed patient cells (MM4, MM5, MM6, MM7 and MM8, Fig.4D-H), Spi alone triggered a pronounced cell killing (Fig.4D-H), with a reduced cell viability of max. up to 60% (Fig.4H). All relapsed patient cells were found to be resistant to Dex-induced cell killing, except MM4, which may explain why a Dex-Spi combination did not further improve on cell killing as compared to Spi alone.

**Table 1:** Patient demographics and disease stage. MGUS, monoclonal gammopathy of undetermined significance; SMM, smoldering myeloma.

| Pseudonym | Gender | Age | Stage | M protein type |
| --- | --- | --- | --- | --- |
| MM1 | F | 60 | Newly diagnosed, prior to first therapy | IgG $\kappa$ |
| MM2 | F | 69 | Newly diagnosed, prior to first therapy | IgG $\kappa$ |
| MM3 | M | 64 | Newly diagnosed, prior to first therapy | $\kappa$ light chain |
| MM4 | F | 71 | Relapsed, prior to start of 6th line therapy | IgG $\lambda$ |
| MM5 | M | 66 | Relapsed, prior to start of 6th line therapy | $\lambda$ light chain |
| MM6 | M | 64 | Relapsed, prior to 2nd line of therapy | IgG $\kappa$ |
| MM7 | F | 71 | Relapsed, prior to 2nd line of therapy | $\kappa$ light chain |
| MM8 | M | 56 | Relapsed, prior to 2nd line of therapy | IgA $\kappa$ |
| MM9 | F | 31 | High risk SMM | IgG $\lambda$ |
| MM10 | M | 53 | MGUS | IgM $\kappa$ |

**Fig. 4:**
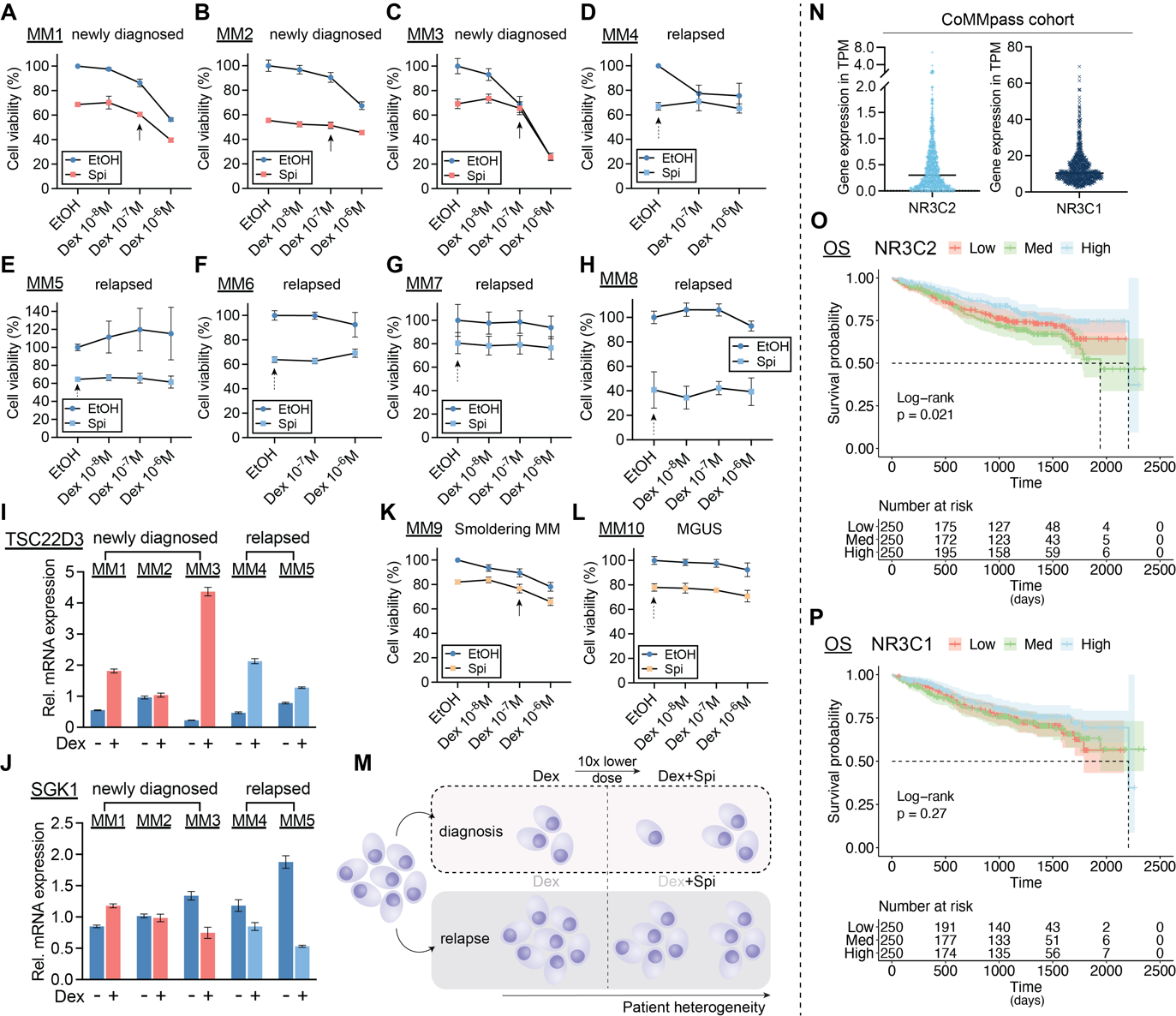
Combining lower doses of Dex with the MR antagonist Spi enhances cell killing in primary myeloma cells depending on the disease stage. (**A-H**) Patient-derived MM cells from bone marrow aspirates of (**A-C**) newly diagnosed, (**D-H**) relapsed or MM patients were treated with a Dex concentration range (10^−6^M-10^−8^M), Spi (10^−5^M), a Dex-Spi combination or solvent control (EtOH) for 24h (**A, C-G**) or 72h (**B, H**), followed by a CelltiterGlo cell viability assay. (**I-J**) When the primary cell yield was sufficient, primary MM cells were treated for 6h with Dex (10^−6^M) or solvent control, followed by RNA isolation and RT-qPCR analyses to determine the expression levels of *TSCD22D3* (GILZ) and *SGK1*. Data analyses were performed using qBaseplus with *SDHA*, *RPL13A* and *YWHAZ* serving as reference genes. The bar plots represent the mean +/− SD of 3 technical replicates. Overall, no statistical analyses were performed because only 1 biological replicate could be carried out given the limited culturing time of primary MM cells isolated from a BM aspirate. (**K-L**) Patient-derived MM cells from bone marrow aspirates of premalignant (smoldering MM or MGUS) myeloma patients were treated with a Dex concentration range (10^−6^M-10^−8^M), Spi (10^−5^M), a Dex-Spi combination or solvent control (EtOH) for 24h (**L**) or 48h (**K**) followed by a CelltiterGlo cell viability assay. (**M**) Graphical summary demonstrating that primary MM cells isolated at diagnosis undergo profound Dex-mediated cell killing, while the addition of Spi to a 10-fold lower Dex dose triggers more extensive cell killing, although not the same extent in all patients. In the relapsed setting, Dex is unable to induce significant primary MM cell killing, while Spi triggers a substantial MM cell killing response. The extent of the described cell killing effects varies from patient to patient, due to interpatient heterogeneity, which is well known in MM. (**N**) TPM (transcripts per million) gene expression values, generated via RNA-sequencing, of *NR3C2* and *NR3C1* in the CoMMpass cohort; only samples at diagnosis were taken along. (**O-P**) Kaplan-Meier curve of the MMRF patient cohort, depicting the survival probability in function of overall survival (OS) for low, medium or high expression of (**O**) *NR3C2* or (**P**) *NR3C1*. Statistical analyses were performed in R (package survival), using a log-rank test. Data information: (**A-H**, **K-L**) Each data point represents the mean +/− SD of technical replicates because only one biological repetition could be performed with the primary myeloma cells. The solvent condition was set as 100% and the other conditions were recalculated accordingly. Full arrows highlight the effect of the combination of a 10-fold lower Dex dose with Spi, while dashed arrows indicate the effect of Spi alone.

Only for MM1, MM2, MM3, MM4 and MM5 patients the primary cell yield was sufficiently high to allow for an analysis of GR and MR target gene expression following Dex treatment. We selected *TSC22D3* and *SGK1* because 1) of their opposing Dex response in our MM cell lines (Supplementary Fig.S1), 2) studies indicate anti-proliferative actions (*TSC22D3*)^37^ or pro-survival effects (*SGK1*)^39^, and 3) the receptors themselves were below the detection limit as assessed by RT-qPCR. Dex treatment upregulated TSC22D3 mRNA levels in 2 out of 3 newly diagnosed patient cells (MM1 and MM3) and in both relapsed patient cells (MM4 and MM5) (Fig.4I), while SGK1 mRNA levels were downregulated in 1 out of 3 newly diagnosed patient cells (MM3) and in both relapsed patient cells (MM4 and MM5) (Fig.4J); hereby recapitulating the varying degree in Dex-responsiveness that was also retrieved in the MM cell lines (Supplementary Fig.S1).

To examine whether *NR3C2* and/or *NR3C1* expression levels could predict survival, we took advantage of publicly available RNA-sequencing data generated in the framework of the CoMMpass study of the MM research foundation (MMRF). We found that *NR3C2* levels were much lower than those of *NR3C1* at diagnosis (Fig.4N). Nonetheless, *NR3C2* levels were predictive for overall survival (OS) when patients were divided in 3 groups based on high, medium, and low expression of *NR3C2* (Fig.4O). This was not the case when progression free survival (PFS) was assessed (Supplementary Fig.S3A). *NR3C1* expression levels were not predictive of either PFS or OS (Fig.4P, Supplementary Fig.4B).

Taken together, in newly diagnosed and premalignant myeloma patients, a 10-fold lower Dex dose in combination with Spi could be advantageous, although the extent of the therapeutic benefit will differ among patients (Fig.4M, top panel). In the relapsed setting, Dex is barely functional, but Spi alone does induce distinct MM cell killing (Fig.4M, bottom panel). Finally, NR3C2, but not NR3C1 expression levels are associated with OS in patients.

### GR and MR interact at the endogenous level in MM cells

Because crosstalk mechanisms between nuclear receptors can arise from a direct interaction^16^, we examined to which extent and in which direction Dex-Spi steers GR-MR heterodimerization compared to Dex, via two complementary methods. First, we developed a NanoBiT-based quantitative GR-MR heterodimerization assay in HEK293T cells that relies on overexpressed tagged receptors and *in cellulo* reconstitution of a functional NanoLuc luciferase (Fig.5A). In this assay, a signal for GR-MR heterodimerization is only measured when GR coupled to SmBiT and LgBiT coupled to MR interact (Fig.5A). We found that Dex triggered an ∼8-fold induction of GR-MR heterodimerization, which was reduced when combined with Spi to ∼6-fold (Fig.5B-C). In contrast, Spi alone failed to induce GR-MR heterodimerization.

**Fig. 5:**
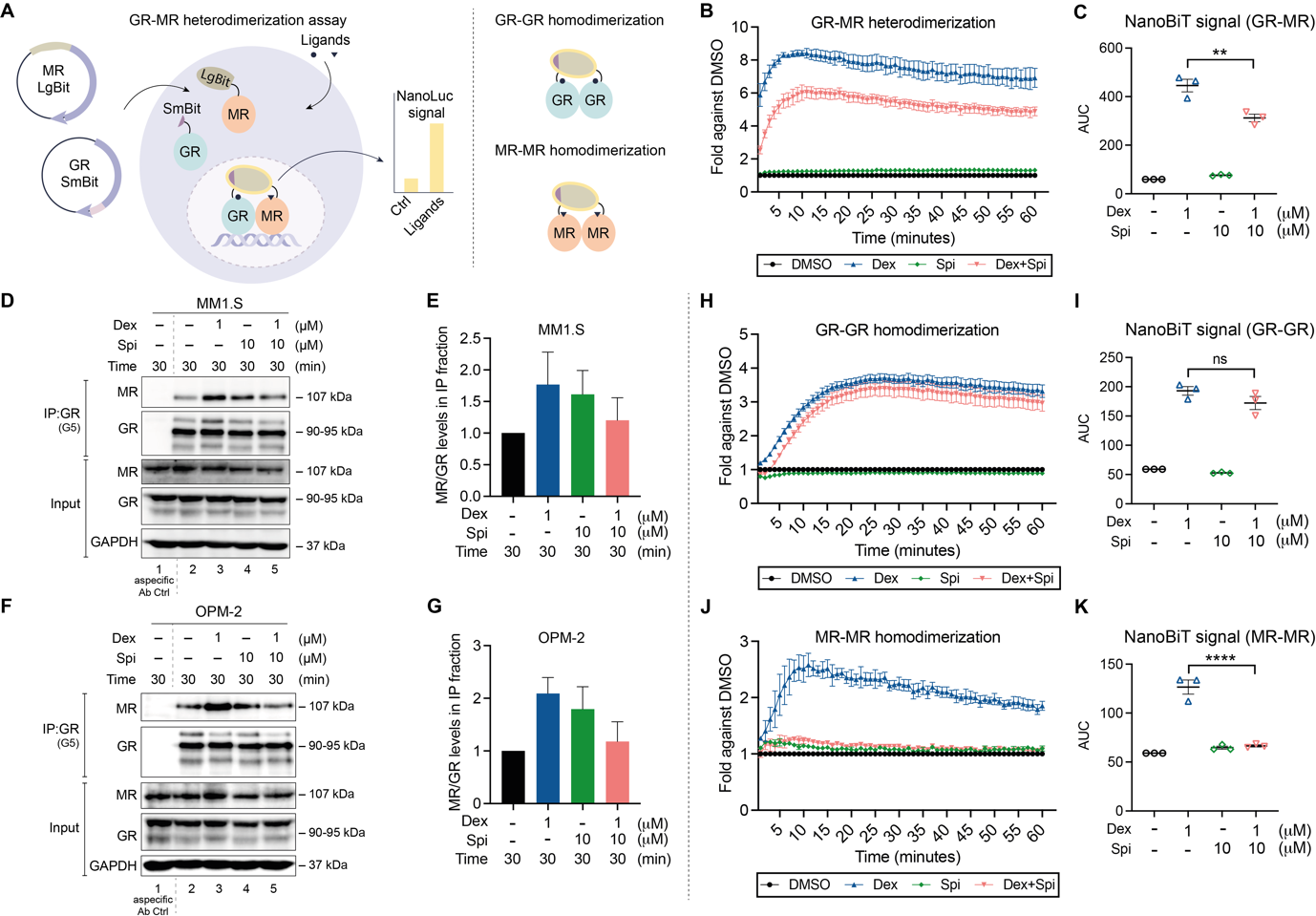
Crosstalk between GR and MR may result from an endogenous interaction that can be modulated with ligands. (**A**) NanoBiT-based GR-MR heterodimerization assay. The Large BiT (LgBiT) and Small BiT (SmBiT) fragments of the NanoLuc® luciferase, which have very low affinity for each other, are coupled to MR (at the N-terminus) or GR (at the C-terminus), respectively, and transfected into HEK293T cells. When the addition of ligand promotes GR-MR heterodimerization, the LgBiT and SmBiT come in close proximity of each other, hereby reconstituting the functional NanoLuc® luciferase. Following substrate addition (furimazine, cell-permeable substrate), the bioluminescent signal can be measured in intact cells. This NanoBiT-based assay was expanded to also measure GR-GR and MR-MR homodimerization. In both cases, LgBiT was coupled to the N-terminus and SmBiT to the C-terminus of both respective receptors. (**B-C**) HEK293T cells were transfected with LgBiT-MR and GR-SmBiT. 24h post-transfection, substrate is added and the baseline luminescence is recorded. Thereafter, cells are treated with Dex (10^−6^M), Spi (10^−5^M), the combination thereof, or solvent control and luminescence is measured continuous during 60min (1min intervals) (N=3). (**C**) Statistical comparison of the area under the curve of Dex vs Dex-Spi NanoBiT results in panel **B** (N=3). (**D-G**) Two myeloma cell lines, i.e. (**D**) MM1.S and (**H**) OPM-2 cells were treated with Dex (10^−6^M), Spi (10^−5^M), a Dex-Spi combination or solvent control for 30min. Protein lysates were prepared and subjected to endogenous immunoprecipitation using GR (G5) antibody (both cell lines N=2). Thereafter, WB analyses were performed to determine co-immunoprecipitation of GR (90-95kDa) with MR (107kDa). GAPDH served as loading control for the input fraction. Lane 1 represents the non-specific antibody control. (**E, G**) In the IP fraction, MR protein levels were quantified relative to GR protein levels by band densitometric analysis using ImageJ. The bar plot displays the ratio of MR/GR in the IP fraction averaged over both biological repetitions (+/ SEM). (**H-K**) HEK293T cells were transfected with (**H**) LgBiT-GR and GR-SmBiT, or (**J**) LgBiT-MR and MR-SmBiT. 24h post-transfection, substrate is added and the baseline luminescence is recorded. Thereafter, cells are treated with Dex (10^−6^M), Spi (10^−5^M), the combination thereof, or solvent control and luminescence is measured continuous during 60min (1min intervals) (N=3). (**I, K**) Statistical comparison of the area under the curve of Dex vs Dex-Spi NanoBiT results in panel **H** and **J** (N=3). Data information: (**D, H**) One representative image is shown for each co-IP experiment; the other biological replicates are available for consultation in Supplementary Fig.5.

We compared the NanoBiT assay results with endogenous GR-MR co-immunoprecipitation (co-IP) analyses in MM1.S and OPM-2 cells. Already in basal conditions, GR and MR interacted in the IP fraction (lane 2, Fig. 5D,F). In line with the NanoBiT results, Dex treatment consistently increased this interaction in MM1.S and OPM-2 cells. Similarly, Dex-Spi combination again reduced this GR-MR interaction compared to Dex treatment. In contrast to NanoBiT, Spi alone did support a marked GR-MR interaction in an endogenous context, although to a lower extent than Dex alone.

To examine whether the Dex-Spi combination could also impact receptor homodimer formation, we extended our NanoBiT assay portfolio towards GR-GR and MR-MR homodimerization. We found that Dex triggered a ∼3.5-fold induction in GR-GR homodimer formation, which was unaffected by the addition of Spi (Fig.5F-G). In contrast, Spi completely abolished the Dex-induced MR-MR homodimer formation (Fig.5J-K).

Summarized, GR and MR engage in an endogenous interaction in MM cells. Quantitative assays indicate that Spi blunts Dex-induced GR-MR and MR-MR dimerization.

### Dex-Spi combination strongly inhibits several major players in myeloma cell survival

To determine whether GR-MR crosstalk leads to differential transcriptomic signatures, we followed an RNA-sequencing approach. Principal component analysis showed that the biological repeats are well clustered per condition (EtOH, Dex, Spi, Dex-Spi) (Supplementary Fig.S6A). Differential gene expression analysis of pairwise comparisons revealed that the highest number of unique genes were regulated by Dex-Spi (596) (Supplementary Fig.6B). Volcano plots further highlight significant genes with the largest log2 fold changes (log2FC; red and blue) for different pairwise comparisons, including ‘Dex-Spi vs EtOH’ (Fig.6A), ‘Dex vs EtOH’ and ‘Spi vs EtOH’ (Supplementary Fig.S6C-D). Several top genes that were shared between pairwise comparisons and that are typical target genes for GR and MR, i.e. *TSC22D3*, *FKBP5* and *SGK1*, were analyzed by RT-qPCR to validate the RNA-sequencing results (Supplementary Fig.S6B). In MM1.S, OPM-2 and L-363 cells, *TSC22D3* (anti-proliferative action)^37^ and *FKBP5* (GR co-chaperone)^40^ mRNA levels were upregulated to a lesser extent by 6h Dex-Spi treatment than by Dex alone, in line with their corresponding count plots (Supplementary Fig.S6D-H). In addition, *SGK1* (stimulates myeloma cell survival)^39^ mRNA levels were downregulated to a similar extent by Dex-Spi and Dex in MM1.S and L-363 cells, again in line with the RNA-sequencing results. In search of significantly regulated side-effect markers, the bone homeostasis marker *TMEM119*^41, 42^ was selected given that this gene may act as a molecular proxy for GC-related bone disease. Dex-Spi combination showed a mild, yet consistent upregulation of *TMEM119* in MM1.S, OPM-2 and L-363 cells (Supplementary Fig.S6D-H).

**Fig. 6:**
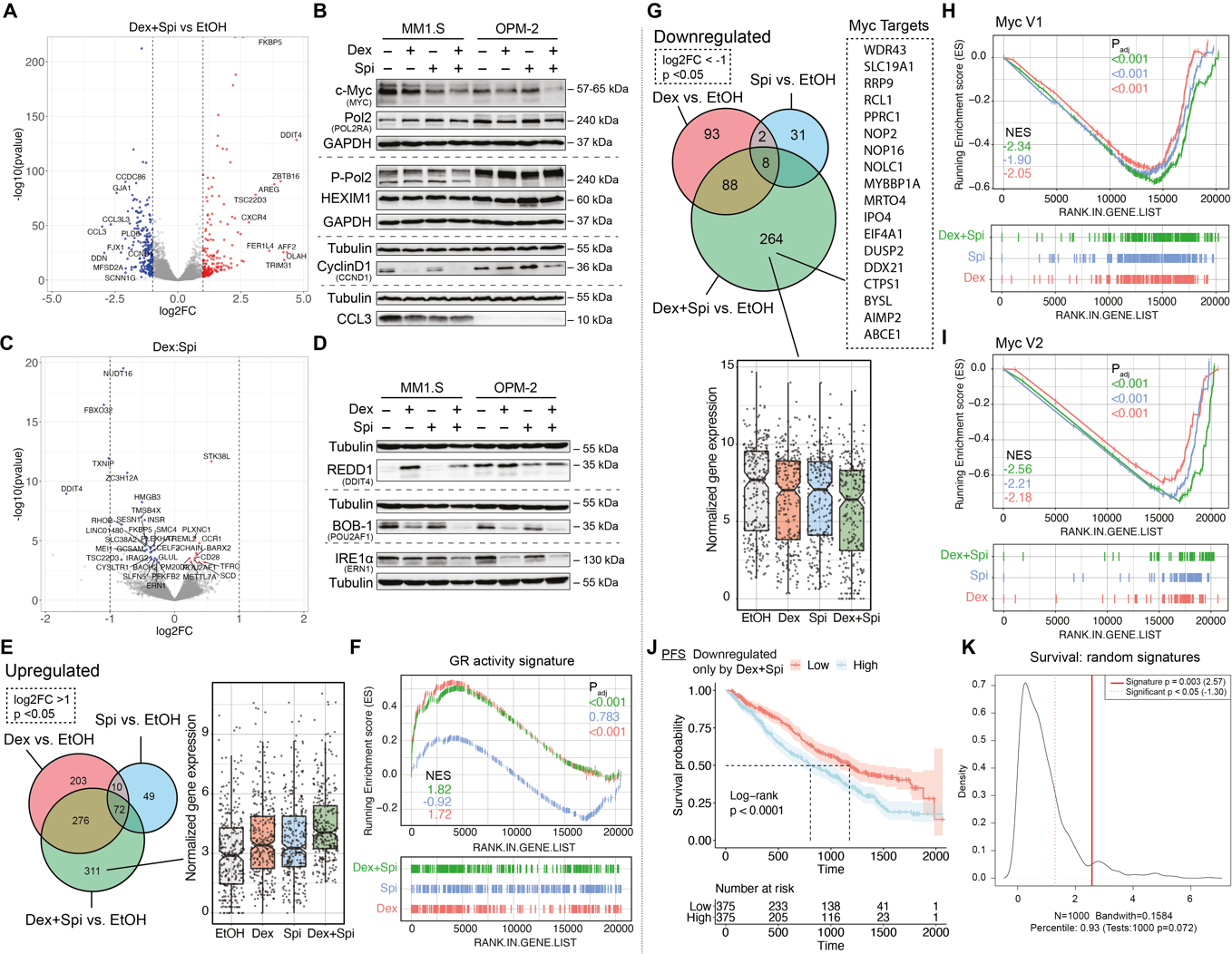
c-Myc and its target genes are inhibited most by Dex-Spi treatment, while a subset of Dex-Spi downregulated genes may predict prognosis. (**A, C**) MM1.S cells were treated with Dex (10^−6^M), Spi (10^−5^M), a Dex-Spi combination or solvent control (EtOH) for 6h, followed by RNA-seq analysis. (**A, C**) Volcano plots depicting the p_adj_ (log10 scale) in function of the log2FC for all genes with baseMean ≥50 for (**A**) the pairwise comparison Dex-Spi vs EtOH or (**C**) the interaction term genes (= those for which the response following Dex-Spi treatment is significantly different from combining the separate responses of Dex and Spi). Significantly regulated genes (p_adj_<0.05) are colored in red (log2FC>1 in **A**, log2FC>0 for **C**, upregulated) or blue (log2FC<-1 in **A**, log2FC<0 for **C**, downregulated); non-significant genes (p_adj_>0.05) in grey. The gene names are displayed for those genes having the largest abs(log2FC) values (top 10 upregulated/downregulated). The dashed lines are set at abs(log2FC)=1. (**B, D**) MM1.S and OPM-2 cells were treated with Dex (10^−6^M), Spi (10^−5^M), a Dex-Spi combination or solvent control (EtOH) for 24h (both N=3). Protein lysates were prepared and subjected to WB analyses, hereby assessing the protein levels of (P-Ser2) RNA-Pol2 (240kDa), GR (90-95kDa), c-myc (57-65kDa), cyclin D1 (36kDa), MIP-1α (CCL3, 10kDa), DDIT4 (35kDa), IRE1α (110-130kDa) and BOB-1 (35kDa). GAPDH (37kDa) and Tubulin (55kDa) served as loading controls. (**E, G**) Venn-diagram of three pairwise comparisons, split up in genes that were either (**E**) upregulated or (**G**) downregulated. In addition, the normalized gene expression profiles of the genes that are uniquely regulated by Dex-Spi are shown. (**F, H, I**) Gene set enrichment analysis (GSEA) of single hallmarks, i.e. (**F**) a GR activity signature and (**H, I**) two sets of Myc target genes (V1, V2), for each pairwise comparison, along with the respective normalized enrichment score (NES) and p_adj_. (**J**) Kaplan-Meier curve of the MMRF patient cohort (N=750), depicting the survival probability in function of progression-free survival (PFS) for low or high expression of genes that were uniquely downregulated by the Dex-Spi combination. Statistical analyses were performed in R (package survival), using a log-rank test. (**K**) Prognostic factor analysis of the genes uniquely downregulated Dex-Spi (red solid curve) versus random signatures (red dotted curve). Prognostic power as determined by SigCheck (R package) of the genes uniquely downregulated by Dex-Spi (red dotted line) with 1000 random gene-sets of the same size (P value <0.05 is indicated by the red dotted line) for the overall survival parameter in the CoMMpass cohort. Data information: (**B, D**) One representative image is shown for each WB experiment, with the number of biological replicates mentioned in each panel description.

To prioritize candidate genes for validation at the protein level, we identified the molecular and cellular functions attributable to the ‘Dex-Spi vs EtOH’ comparison by performing an Ingenuity Pathway Analysis (IPA). We found groups of genes that were significantly involved in gene transcription (terms: gene expression and RNA post-transcriptional modification), cell death and survival and cell cycle (Supplementary Fig.S7A). Based on the top regulated genes from each comparison in these IPA terms (Fig.6A, Supplementary Fig.S6C-D, S7A), we selected genes that were involved in transcriptional regulation (*POLR2A, HEXIM1*), cell growth, proliferation and/or survival (*MYC, CCND1, CCL3*) and validated these at the protein level in MM1.S, OPM-2, L-363 and MM1.R cells (Fig.6B, Supplementary Fig.S7B-C). Interestingly, c-myc (oncogene)^43^ protein levels were significantly decreased by Dex-Spi as compared to Dex alone in both MM1.S and OPM-2 cells (Fig.6B), again in line with our RNA-sequencing results, and also by Spi alone in MM1.R cells (Supplementary Fig.S7C). Whereas RNA polymerase II (Pol II) levels were largely unchanged comparing Dex-Spi versus Dex alone, activated Pol II protein levels, hallmarked by Ser2 phosphorylation^44^, decreased markedly upon Dex-Spi, yet only in MM1.S cells (Fig. 6B); indicating a halt in the transcription elongation process. In OPM-2 cells, a brake on transcription induced by Dex-Spi may rather originate from a mild upregulation (vs Dex) of the transcriptional repressor HEXIM1^45^. Cyclin D1, a protein downregulated by GCs to induce cell cycle arrest^46^, was downregulated to the same extent by Dex-Spi as by Dex alone in MM1.S cells, while in OPM-2 cells, Dex-Spi triggered the largest decrease in cyclin D1 protein levels. In addition, in MM1.S cells, the protein levels of CCL3, a contributor to myeloma cell migration and an aggravator of bone disease^47^, were decreased similarly in all conditions versus solvent (Fig.7B). In OPM-2 and L-363 cells, CCL3 was largely undetectable.

**Fig. 7:**
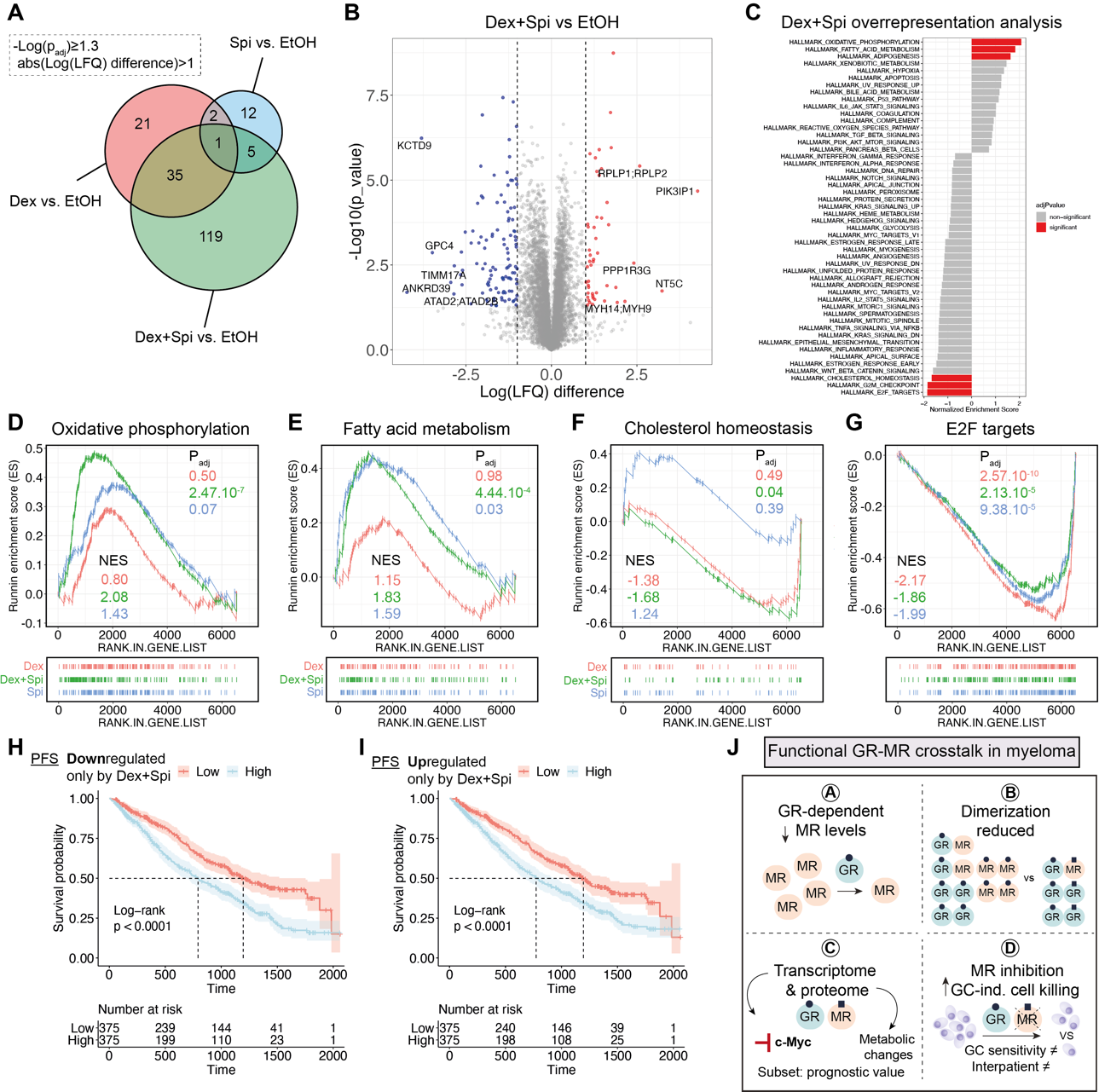
Several metabolic pathways are deregulated most by the Dex-Spi combination treatment. (**A**) MM1.S cells were treated with Dex (10^−6^M), Spi (10^−5^M), a Dex-Spi combination or solvent control (EtOH) for 24h, followed by MS-based shotgun proteomics. Venn-diagram of pairwise comparisons in which significantly regulated proteins (-log(p_adj_)≥1.3) with a abs(log(LFQ difference))>1 were considered. (**B**) Volcano plot depicting the p_adj_ (log10 scale) in function of the log(LFQ) in the pairwise comparison Dex-Spi vs EtOH. Significantly regulated proteins -log(padj)≥1.3 are colored in red (log(LFQ)>1, upregulated) or blue (log(LFQ)<-1, downregulated); non-significant genes (-log(p_adj_)<1.3) in grey. (**C**) GSEA-based overrepresentation analysis for the proteins regulated by Dex-Spi, hereby identifying hallmarks that are significantly (red) or non-significantly (grey) enriched. (**D-G**) GSEA of single hallmarks, i.e. (**D**) oxidative phosphorylation, (**E**) fatty acid metabolism, (**F**) cholesterol homeostasis or (**G**) E2F targets, for each pairwise comparison, along with the respective normalized enrichment score (NES) and p_adj_. (**H-I**) Kaplan-Meier curves of the MMRF patient cohort (N=750), depicting the survival probability in function of progression-free survival (PFS) for low or high expression of proteins that were uniquely (**H**) downregulated or (**I**) upregulated by the Dex and Spi combination. Statistical analyses were performed in R (package survival), using a log-rank test. (**J**) Several lines of evidence support a crosstalk between GR and MR in MM: A) GCs induce a GR-dependent MR downregulation; B) GR and MR engage in a direct, physiologically relevant endogenous interaction that can be modulated by ligands. Spi was shown to reduce the Dex-induced GR-MR heterodimer levels and abolished Dex-induced MR-MR homodimers. Spi did not impact Dex-induced GR-GR homodimerization; C) Dex and Spi combination gives rise to a differential gene and protein expression profile, in which the inhibition of c-myc and its target genes, and several metabolic pathways are modulated most pronounced by Dex-Spi, respectively. A specific subset of targets may even have prognostic significance; D) MR inhibition enhances GC-induced cell killing in MM cell lines depending on their GC responsiveness and in primary (heterogeneous) MM cells depending on the disease stage.

We also examined the Dex:Spi interaction term (Fig.6C), which contained 37 significantly regulated genes (Supplementary Fig.S1) and of which the response following Dex-Spi treatment is hypothesized to be significantly different from combining the responses of separate Dex and Spi treatments. Three genes were selected for validation at the protein level based on their high normalized counts, previously described function in myeloma, and accompanying primary antibody performance (in WB): 1) *DDIT4* (also known as REDD-1), promotes myeloma cell growth and survival^48^ and is described as a GC-inducible muscle atrophy marker^49^; 2) *ERN1* (also known as IRE1α), a sensor of unfolded proteins and critical for MM tumor growth^50^; 3) *POU2AF1* (also known as BOB-1), a regulator of oncogenic networks in myeloma^51^. Interestingly, REDD1 protein levels were decreased by Dex-Spi combination versus Dex in MM1.S cells and, to a similar extent, by Spi and Dex-Spi in OPM-2 cells (Fig.6D). In L-363 cells, REDD1 levels were equally increased by Dex and Dex-Spi (Supplementary Fig.S7D). Both BOB-1 and IRE1α were most strongly decreased by Dex-Spi combination in MM1.S cells and comparably decreased by Dex and Dex-Spi treatment in OPM-2 cells (Fig.6D). In L-363 and MM1.R cells, IRE1α levels were not clearly regulated by any treatment (Supplementary Fig.S7D).

Taken together, our transcriptome analysis and subsequent validation at the protein level reveals that gene transcription is halted more strongly by Dex-Spi than by Dex alone in MM cells, with c-myc, cyclinD1, REDD1 and BOB-1 being strongly Dex-Spi-downregulated targets.

### c-myc target genes are mostly downregulated by Dex-Spi

We expanded our RNA-seq analysis by zooming in on genes that were uniquely up-or downregulated by the Dex-Spi combination. We found 311 genes to be uniquely upregulated by Dex-Spi, as shown by their normalized gene expression profile (Fig.6E). GSEA shows that genes upregulated by Dex-Spi (green curve) or Dex (red curve) treatment were enriched for a previously established GR activity score^52^ (Fig.6F).

We then focused on the 264 genes that are uniquely downregulated by Dex-Spi treatment and for which their normalized gene expression is depicted in Fig.6G. GSEA-based overrepresentation analysis (Supplementary Fig.S7E) identified that two hallmarks containing different sets of myc target genes were significantly enriched in the set of genes that are uniquely downregulated by Dex-Spi. Zooming in on these ‘myc hallmarks’ (Fig.6H-I) confirmed that the most negative normalized enrichment scores (NES) was indeed obtained for Dex-Spi. In addition, we examined whether the expression levels of the uniquely Dex-Spi downregulated genes could predict survival, for which we again relied on the CoMMpass cohort data. Strikingly, we found that patients showing low expression of the Dex-Spi downregulated genes had better PFS and OS compared to patients showing high expression of these genes (Fig.6J, Supplementary Fig.S7F). Prognostic factor analysis highlighted that this prediction is better than when random gene signatures were used (Fig.6K, full red line).

Overall, the inhibition of c-myc target genes likely underpins the enhanced myeloma cell killing observed with Dex-Spi.

### Shotgun proteomics analyses unveils a contribution of metabolic pathway deregulation upon Dex-Spi treatment

To gain additional mechanistic insights into the Dex-Spi combination treatment at 24h treatment, we performed mass spectrometry-based shotgun proteomics on MM1.S cells. Differential expression analysis showed that the highest number of hits (i.e. proteins) was again identified for the Dex-Spi combination treatment (Fig.7A); in line with RNA-sequencing results (Supplementary Fig.S6B). Volcano plots further highlighted significant proteins with the largest log(LFQ) difference (red and blue) for different pairwise comparisons, including ‘Dex-Spi vs EtOH’ (Fig.7B), ‘Dex vs EtOH’ and ‘Spi vs EtOH’ (Supplementary Fig.S8A,C). Overrepresentation analyses of the Dex-Spi regulated proteins identified that several hallmarks of metabolism, including oxidative phosphorylation and fatty acid metabolism were significantly upregulated, while the hallmarks for cholesterol homeostasis, G2M checkpoint and E2F targets were significantly downregulated (Fig.7C). Zooming on the individual hallmarks (Fig.7D-G, Supplementary Fig.S8E-H) learned that only proteins regulated by Dex-Spi, and not by Dex or Spi, were significantly enriched for oxidative phosphorylation and fatty acid oxidation (Fig.7D-E). Cholesterol homeostasis was rather inhibited by the Dex-Spi regulated proteins (Fig.7F). In contrast, all treatments (Dex-Spi, Dex and Spi) gave rise to a significant enrichment of the G2M checkpoint and E2F targets, but it was rather Dex alone that inhibited these hallmarks the most (Fig.7G, Supplementary Fig.7E). In addition, the Myc V2 hallmark was only significantly enriched upon Spi treatment in the proteomics analyses (Fig.S8G-H), while this enrichment was significant for all treatments in the RNA-sequencing analysis (Fig.6H-I), which may be due to a difference in treatment time in both setups (6h vs 24h). Nonetheless, the master regulator c-myc was differentially regulated at the protein level (Fig.6B).

Finally, we examined whether the expression levels of Dex-Spi regulated proteins could predict survival in the CoMMpass cohort. The genes corresponding to the proteins that were uniquely up-or downregulated by Dex-Spi (upset plots, Supplementary Fig.8E-F) were used as input as only transcriptomics data are available for the CoMMpass cohort. We found that patients having a low expression of the proteins uniquely downregulated by Dex-Spi had a better PFS and OS than patients having a high expression of these targets (Fig.7H, Supplementary Fig.S9A). In contrast, high expression levels of proteins that were uniquely upregulated by Dex-Spi identified patients with worse PFS and OS (Fig.7I, Supplementary Fig.S9B).

Summarized, on a mechanistic level, we additionally resolved that the Dex-Spi combination treatment deregulates several metabolic pathways.

## Discussion

In this study, we have identified a novel, functionally relevant nuclear receptor crosstalk mechanism between GR and MR in myeloma cells (summarized in Fig.7J). We have shown that although MR levels were decreased upon Dex treatment over time in a GR-dependent manner (A; Fig.7J), endogenous GR strongly interacts with MR in a Dex-inducible manner at early stages (B; Fig.7J). We further found that Spi clearly diminished Dex-induced GR-MR heterodimerization and completely abolished Dex-induced MR-MR homodimerization (B, Fig.7J). Dex-Spi combination treatment also gave rise to a differential transcriptomic and proteomic signature (C; Fig.7J) that can help explain the enhanced Dex-induced myeloma cell killing in combination with Spi (D; Fig.7J). These four main findings will be discussed in further detail below.

We found that Dex downregulates MR levels likely by a superposition of different mechanisms. In earlier work, GCs were reported to reduce the stability of pro-inflammatory mediators, such as TNFα, by upregulating mediators of mRNA decay^53^. A similar mechanism decreased the *NR3C2* mRNA stability in renal epithelial cells subjected to hypertonic conditions^54^, and recently several *NR3C2*-targeting miRNA’s were identified in this context^55^. In myeloma cells, Dex did not further reduce the MR mRNA stability following ActD treatment (Fig.1I). Pharmacological inhibition of transcription and translation rather supported that the Dex-induced decline of MR mRNA and protein depended, at least partially, on both mechanisms. Noteworthy, Dex treatment also induced a second MR band, approximately 10kDa upwards, hinting at various post-translational modifications. Although an Aldosterone-induced upward shift of MR of even 30kDa was reported before, this was linked to increased phosphorylation on several serine residues with subsequent MR polyubiquitination and proteasomal degradation^56^. In the MM cell context however, Dex did not decrease MR protein levels via proteasomal or lysosomal degradation (Supplementary Fig.S2B-E). Although GR was clearly required for the Dex-induced MR downregulation (Fig.1B,F,G), an additional regulatory mechanism whereby GCs may directly bind to and affect MR, as in MM1.R cells that lack GR (Supplementary Fig.S2A), cannot be excluded.

We are to the best of our knowledge the first to report that endogenous GR and MR may form heterodimers or are at least part of the same protein complex in myeloma cells. Compared to typical nuclear receptor heterodimers (e.g. RXR and PPAR), atypical heterodimers such as GR-MR could be less prominent and transient, but that does not exclude a potentially strong functional effect^16^. Although GR and MR connect already in basal conditions, their interaction was further supported by Dex via two orthogonal assay systems (Fig.5B-G). Spi, however, consistently reduced Dex-induced GR-MR heterodimerization (Fig.5B-G) and completely abolished MR-MR dimerization (Fig.5J-K). Together, these results support a hypothesis that altered receptor dimerization equilibria may mechanistically contribute to an altered transcriptome and proteome profile and ultimately to the enhanced cell killing induced by the Dex-Spi combination. Besides direct, also indirect crosstalk mechanisms can affect the therapy response, as shown for GR and AR in prostate cancer, where even diminished responsiveness to enzalutamide (anti-androgen) was observed^17, 57^. Although the GR-MR crosstalk in MM1.S and OPM-2 cells may entail a direct physical interaction (Fig.5B-E,5H-I), further studies are necessary to discriminate between tethering-based interactions or cooperative DNA binding modes^23, 27^ in the context of myeloma.

Transcriptome analysis has shown that Dex-Spi halted markers of transcription elongation. We found decreased Pol II Ser2 phosphorylation in MM1.S cells (Fig.6B), which agrees with earlier work resolving tethering-based GR repression mechanisms in an inflammatory setting. There, Dex-activated GR hampered Pol II Ser2 phosphorylation at several NF-κB-regulated promoters^44^. In OPM-2 cells, increased expression of the transcriptional repressor HEXIM1 appeared more decisive for a Dex-Spi-induced block in transcription elongation (Fig.6B). The latter findings agree with a study where HEXIM1 sequestered positive transcription elongation factor b (P-TEFb) to inhibit transcription elongation of tumorigenic genes^45^. In line, Rogatsky and colleagues demonstrated that GR can even inhibit recruitment of the P-TEFb complex that is normally responsible for Pol II Ser2 phosphorylation^58^. Altogether, Dex-Spi consistently triggers several (consecutive) steps to inhibit transcription in GC-sensitive MM cells, regardless of cell-line specific regulations of the underlying markers.

Furthermore, our results strongly suggest that c-myc and many of its target genes (Fig.6B, 5H-I) may be responsible for the enhanced myeloma cell killing induced by Dex-Spi. The fact that GCs decrease c-myc levels is well documented in literature^46, 59^, and results in cell cycle arrest at the G1 phase in leukemia cells^59^; even Spi alone was linked to decreased c-myc activity before^60^. In myeloma, reports show that c-myc protein was overexpressed in 40% of patients at diagnosis, which correlated with shorter OS^43^. Moreover, in 2022, the team of Rosen showed that inhibiting SUMOylation in myeloma cells resulted in decreased c-myc protein stability, which in turn decreased the levels of several miRNAs involved in either GR downregulation or GC resistance^61^. Using survival analyses on the CoMMpass patient cohort (Fig.6J-K), we further found that patients have a lower risk of progression when displaying low levels of the unique Dex-Spi-downregulated genes, which may altogether be predictive of the clinical relevance of a combination treatment. In addition, patients having high MR expression levels at diagnosis showed superior OS, while GR expression levels were not predictive for survival (Fig.4O-P). For GR, this contrasts a previous study, where high expression levels at diagnosis were found predictive for OS^62^. One reason for this difference may be that Rosen and colleagues stratified patients in two subgroups, based on whether they underwent stem cell transplantation or not, and another reason may be that at that time only version IA13 of the database was available (with 650 patients vs. IA14 with 750 patients).

Our study of the proteome additionally revealed that metabolic hallmarks such as oxidative phosphorylation and fatty acid metabolism were upregulated, while the hallmark cholesterol homeostasis was downregulated most by the Dex-Spi combination treatment (Fig. 7D-E). Reports in several lymphoid malignant cell types have associated enhanced metabolism, i.e. increased glycolysis, oxidative phosphorylation, cholesterol biosynthesis and fatty acid oxidation, with decreased GC responsiveness or even GC resistance^63–65^. GCs inhibit glycolysis in ALL cells, which is not sufficient to trigger cell death but does induce a metabolic shift to mitochondrial oxidative phosphorylation to obtain survival energy^66^. In line with this, combining GCs with the oxidative phosphorylation inhibitor oligomycin sensitized GC-resistant ALL cells to cell killing^64^. This team also found synergistic cell killing in GC resistant ALL cell lines when GCs were combined with an inhibitor of cholesterol metabolism (simvastatin)^64^. In CLL cells, GCs reduce metabolic activity among others by downregulating pyruvate kinase M2 and decreasing levels of pyruvate. Concomitantly however, this elevated the dependency of the CLL cells on fatty acid oxidation because GCs also upregulated PPARα and PDK4 expression^65^. Based on these studies, increased oxidative phosphorylation and fatty acid metabolism may be rather an unwanted feature of the Dex-Spi combination treatment in a context of prolonged treatment, although this requires further investigation. Within this context, our survival analysis (Fig.7I) supports that high expression of proteins that were uniquely upregulated by Dex-Spi was indeed associated with worse PFS. In contrast, the marked inhibition of cholesterol homeostasis observed solely with Dex-Spi treatment may rather be a contributing mechanism that can drive and explain the enhanced myeloma cell killing. Nonetheless, the connection between GCs and potential shifts in metabolism in myeloma cells, especially in the context of prolonged treatment, is rather understudied, which opens opportunities for follow-up research.

Concerning markers mimicking GC-related side effects, we have found that Dex-Spi increases *TMEM119* mRNA levels in all myeloma cell lines, except in MM1.R (Supplementary Fig.S6D-H). Because this gene supports osteoblast differentiation and bone formation^67, 68^, this suggests that Dex-Spi may improve GC-induced osteoporosis. We have also shown that REDD1 (*DDIT4*), an instigator of myeloma cell growth and survival^48^, is decreased by Dex-Spi combination compared to Dex in MM1.S and OPM-2 (Fig.6D). Because GC-mediated increases in REDD1 levels were also shown to contribute to muscle atrophy^69^, this suggests that Dex-Spi may perhaps improve GC-induced muscle atrophy. Whether those and other metabolic side effects could also be improved at the organism level remains to be investigated in follow-up studies.

We discovered that Spi enhances Dex-induced cell killing of myeloma cells, i.e. in GC-sensitive MM1.S and OPM-2 cells as well as in patient cells of several newly diagnosed patients and a smoldering MM patient (Fig.2, Fig.3A, Fig.4A,B,K). Our findings agree with a study in which a GC-treated pre-B lymphoma cell line stably overexpressing the N-terminal domain (NTD) of MR resulted in blocked apoptosis^70^. In our case, Spi addition partially suppressed Dex-induced GR-MR heterodimer formation and abolished Dex-induced MR-MR homodimerization, which may form a molecular basis to support differential gene and protein expression profiles with enhanced anti-myeloma outcomes compared to Dex. Noteworthy, Spi is a potent FDA-approved MR antagonist, however, less selective because it causes anti-androgenic (via AR) and progestogenic (via progesterone receptor, PR) side effects as well as hyperkalemia^20^. A limitation of our study is that we did not include a more selective MR antagonist, such as Eplerenone, mainly because this compound has a 40-fold lower affinity for MR than Spi^20^. Nonetheless, Spi did not require GR for its action, because GR-negative MM1.R cells underwent ∼30% cell killing upon Spi treatment (Fig.3D). Data show that Spi by itself can suppress pro-inflammatory cytokines and induce apoptosis in blood mononuclear cells by reducing NF-κB and c-myc activities^60^. Spi was also found to inhibit nucleotide excision repair which resulted in increased sensitivity of (primary) myeloma cells to alkylating agents such as melphalan^71^. Our study further supports a marked reduction in cell viability observed upon 10^−5^M Spi monotherapy across all (patient-derived) MM cells (Fig.2-4).

In conclusion, our results support the high potential of MR as an additional therapeutic target in myeloma, of which antagonists may be repurposed for myeloma treatment in combination with GCs as add-on to the myeloma standard of care treatment. We showed that a functional crosstalk between GR and MR exists in myeloma and that a targeting hereof with ligands warrants further investigation of its potential therapeutic benefit in terms of efficacy, safety and the possibility to reduce the GC-dose.

## Materials and Methods

### Cell lines and reagents

MM1.S, OPM-2, L-363, U-266 and MM1.R cells were cultured in RPMI1640 GlutaMAX and HEK293T and EA.hy926 cells in DMEM, both supplemented with 10% fetal bovine serum (FBS), 100U/mL penicillin and 0.1mg/mL streptomycin and grown at 5% CO_2_ and 37°C. MM1.S, MM1.R and EA.hy926 were purchased from ATCC. OPM-2 were kindly provided by Prof. B. Thompson (University of Texas Medical Branch) and L-363 and U-266 cells by Prof. M. Engelhardt (Uniklinik Freiburg, Germany). HEK293T were obtained from the cytokine receptor lab (Ghent University). All cell lines were mycoplasma negative (MycoAlert kit, Lonza). Experiments were performed using charcoal-stripped serum (CTS), unless otherwise specified.

Total solvent concentrations were equal in all conditions. Dex, Hydrocortisone (Hcort), Prednisolone (Pred), fluocinolone acetonide (FA), Aldosterone (Ald), RU486 and cycloheximide (CHX) were purchased from Sigma Aldrich and dissolved in ethanol (EtOH), unless otherwise specified. Spi and Chloroquine (CQ) were obtained from Santa Cruz Biotechnology and dissolved in respectively EtOH and water, unless otherwise specified. MG132 was purchased from Selleck Chemicals and dissolved in DMSO. Flag-GR has been described before^72^ and the GFP-MR construct was a kind gift from Dr. H. Tanaka (University of Tokyo, Japan).

### siRNA nucleofection

MM1.S cells were transfected with siCtrl, siGR or siMR (see Supplementary Table S2) in 24-well plates by nucleofection using cell line nucleofector kit V and the nucleofector device at program X01. 48h post-nucleofection, cells were reseeded to 96-well plates and treated for another 24h with compounds (details in Fig. legends).

### NanoBiT-based homo-and heterodimerization assays

HEK293T cells were seeded in 96-well plates in 10%FBS DMEM and transfected 24h later with 2.5ng pLgBiT-MR and 2.5ng pGR-SmBiT (GR-MR heterodimerization assay) or 1.5ng pLgBiT-GR and 1.5ng pGR-SmBiT (GR-GR homodimerization assay) or 1ng pLgBiT-MR and 1ng pMR-SmBiT (MR-MR homodimerization assay) using calcium phosphate precipitation.

24h later, the Nano-Glo® Live Cell reagent was reconstituted (Promega) and 25μL was added to the transfected cells, after which the baseline luminescence was measured for 15min (continuous mode, 1min intervals) using an Envision (Perkin Elmer) spectrophotometer. Subsequently, ligands were added (see Fig. legends) and the luminescence was measured in a time window of 60min (continuous mode, 1min intervals). Luminescence counts were normalized to baseline and set as a fold-difference versus the solvent condition (here: DMSO). The area under the curve method was used to statistically compare Dex and Dex-Spi conditions.

### RT-qPCR

Total RNA was isolated using the RNeasy mini kit (Qiagen). Reverse transcription (RT) was performed using the iScript cDNA synthesis kit (Bio-Rad). The resulting cDNA served as template for the quantitative PCR (qPCR) reaction, for which Lightcycler 480 SYBR Green I Master mix (Roche diagnostics) was used. Primer sequences are available in Supplementary Table S3. Cq values were analyzed using qBasePlus (Biogazelle) and normalized to the reference genes SDHA, RPL13A and YWHAZ.

### RNA-sequencing

Total RNA was isolated using the RNeasy mini kit (Qiagen). The RNA-seq library was prepared using the Illumina TruSeq stranded mRNA library kit, followed by single-end 100 bp sequencing on a Illumina NOVASeq 600 instrument (VIB Nucleomics core), yielding 19-27 million reads per sample. Briefly, sequencing reads were quality controlled with FastQC (version 0.11.9) and trimmed using Trim-Galore (version 0.6.6-0) to remove low-quality ends (phred score <30) as well as adapters, followed by another quality control of the trimmed data. Thereafter, reads were pre-mapped to PhiX genome using STAR (version 2.7.6a) and the resulting PhiX-unmapped reads were aligned to the human genome GRCh38. The position-sorted output BAM files were converted to count data using HTSeq (version 0.12.4) in the ‘union’ mode. Differential gene expression analysis was performing using DESeq2 R package (version 1.34.0), using an interaction model (design formula: c_0_x_0_ + c_1_x_1_ + c_2_x_2_ + c_3_x_1_x_2_). As input for the analysis, only genes with counts > 1 were withheld. Normalized counts were either plotted per gene or were compared for all genes, clustered, and presented as heatmaps (pheatmap package, version 1.0.12). Pairwise comparisons between differentially treated samples (e.g. Dex-Spi vs EtOH) as well as the interaction term were retrieved at a significance level of α= 0.05, corresponding to Wald-test adjusted p-value (FDR) cutoff (p_adj_). Volcano plots were made depicting the p_adj_ (log10 scale) in function of the log2FC for all genes with baseMean ≥50 in the interaction term and each pairwise comparison of interest. Functional annotations of differentially expressed genes were performed using Ingenuity Pathway Analysis (IPA) or gene-set enrichment analysis (GSEA, using standard parameters)^73^.

### Protein lysates and Western blotting (WB)

Protein lysates were prepared using Totex lysis buffer, as described before^36^, loaded on an SDS-PAGE gel, and blotted onto nitrocellulose membranes (Bio-Rad). The list of primary antibodies can be found in Supplementary Table S4. Note that the primary MR antibody is of a non-commercial source and hence different batches were used throughout the course of this research (clone 6G1, kind gift Dr. Gomez-Sanchez). As secondary antibodies, we used species-specific HRP-conjugated antibodies (cat nr: NA931, NA934, GE-Healthcare). To visualize results, Pierce ECL (Plus) (Thermo Fisher Scientific), Westernbright Quantum or Sirus (Isogen), or ECL Prime (GE Healthcare) served as chemiluminescent substrates and signals were developed using X-Ray films or imaged on a ProXima 2850 (Isogen) or Amersham 680 (GE healthcare) imaging system. Band densitometric analyses were performed using ImageJ.

### Shotgun proteomics

MM1.S were treated for 24h with compounds (see Fig. legends), after which the cells were collected by washing with ice-cold PBS and storing the cell pellets at −80°C. Four biological replicates were performed. The mass spectrometry sample preparation and computational analysis were performed as previously described^52^.

### Co-immunoprecipitation

Post-treatment, MM1.S or OPM-2 cells were lysed in NP-40 lysis buffer (50mM Tris-HCl pH 8.0, 150mM NaCl, 1% NP-40) and subjected to immunoprecipitation using anti-GR G5 antibody, as described before^74^. Briefly, cell lysates were precleared with immobilized protein A dynabeads (50μL bead slurry, with f.c. 2mg/mL BSA) by 1h rotation at 4°C. Ensuing, 100-150μg total protein was combined with anti-GR G5 antibody (sc-393232) and rotated for 1h at 4°C, after which immobilized dynabeads (50μL bead slurry, with f.c. 2mg/mL BSA) were added followed by another 2h rotation at 4°C. Following washing steps, the bead-mixtures were denatured for 5min at 95°C using 4xLaemli buffer supplemented with DTT (f.c. 200mM). Samples were subjected to WB analyses and anti-MR 6G1 antibody (kind gift Dr. Gomez-Sanchez) was used to assay the interaction between immunoprecipitated GR and MR.

### Flow cytometry

MM1.S cells were resuspended in Annexin-binding buffer and between 10^5^ and 5×10^5^ cells were stained with Alexa Fluor 488 Annexin V and propidium iodide (Molecular Probes by Invitrogen). Unstained and single stained cells served as controls. Samples were measured on an Attune Nxt flow cytometer (Thermo Fisher Scientific). Data analysis was performed using FlowJo; the gating strategy is depicted in Supplementary Fig.S3b.

### Cell viability assays

MM cells were seeded and treated immediately with compounds for 24h or 72h (see Fig. legends). Thereafter, cells were subjected to a CellTiterGlo cell viability assay (Promega), as described before^36^. Briefly, the reconstituted CellTiterGlo reagent (Promega) was added in a 1:1 ratio to the cells, and contents were mixed for 2min on an orbital sharker. Following signal stabilization (10min), luminescence was recorded using a Spectramax Paradigm (Beckman Coulter), Envision or Ensight (Perkin Elmer) spectrophotometer.

### Patient-derived MM cells

Sample acquisition was approved by the ethical commission of the Ghent University Hospital (EC UZG 2018/0906) and informed consent was obtained from all patients. Bone marrow aspirates were filtered through a cell strainer and mixed with a RosetteSep human MM cell enrichment cocktail (negative selection, STEMCELL Technologies). Afterwards, bone marrow aspirates were diluted 1:1 with PBS (+ 2% FBS) and layered on a Lymphoprep gradient using SepMate tubes (STEMCELL Technologies). After centrifugation, the cells were washed twice with PBS (+ 2%FBS) and with a red blood cell lysis buffer (0.8% NH_4_Cl, 0.1mM EDTA, STEMCELL technologies). Thereafter, the enriched MM cells were resuspended in RPMI1640 GlutaMAX (+10%CTS) and subjected to a cell viability assay and/or RNA isolation.

### Survival analysis

A publicly available dataset was used to evaluate the prognostic significance of set of genes/proteins identified via RNA-sequencing or shotgun proteomics. Specifically, the Relating Clinical Outcomes in MM to Personal Assessment of Genetic Profile (CoMMpass) trial release IA14 was used, launched by the MM research foundation (MMRF). Normalized TPM gene expression values, generated using RNA-sequencing, were downloaded alongside clinical data through the MMRF research portal (https://research.themmrf.org). Overall survival (OS) was defined as the time from diagnosis until death from any cause or until the time point the patient was last known to be alive. In the latter case patients were censored. Progression free survival (PFS) delineates the time from treatment initiation until relapse or death from any cause. Patients were divided in 2 or 3 groups based on the average of their z-score normalized expression data, ranked from low to high (2 groups) or from low to medium to high (3 groups). Survival analysis of the CoMMpass cohort was performed using using R (package survival, V3.5-3); statistical significance was calculated using the log-rank test. Prognostic factor analysis was done using SigCheck package (V2.28.0), running with standard parameters.

### Statistical analyses

Statistical analyses were performed using GraphPad Prism 9 or R, as specified in the figure legends. Sample size calculations were not performed upfront. Experiments were performed in at least three independent repetitions, as detailed in the figure legends, except for experiments involving patient material, which could only be performed once because of the limited culturing time and yield of the isolated primary cells. Error bars represent the standard error of the mean (SEM), except for experiments involving primary patient material, where the error bars represent the standard deviation (SD). When the means of 2 groups were compared a two-tailed independent Student’s t-test was used, when the means of more than 2 groups were compared a one-way or two-way ANOVA with (Tukey’s or Sidak’s) multiple comparisons post-test was used, as detailed in the figure legends. Normal distribution and equality of variances were assumed. Statistical significance in survival curve estimates were calculated using the log-rank test. When *P* < 0.05, results were designated significant: * = *P* < 0.05, ** = *P* < 0.01, *** = *P* < 0.001, **** = *P* < 0.0001, ns = non-significant.

## Data availability

The datasets used in this study are available in the following databases: RNA-seq data in the Gene Expression Omnibus (GEO) database with accession number GSE200313; shotgun proteomics data are scheduled for deposit to the Proteomics Identification (PRIDE) database.

## Acknowledgements

DC was funded by a predoctoral fellowship from Flanders Innovation and Entrepreneurship (VLAIO), grant number 131374 (www.vlaio.be) until December 2017. Research project funded by Kom op tegen Kanker (Stand up to Cancer), the Flemish cancer society (grant numbers KDB 2012-VLK-GC-MM and STI.VLK.2018.0019.01). This work was further supported through VIB and Ghent University institutional funding to KDB.

## Author contributions

**Dorien Clarisse**: Conceptualization; investigation; data curation; formal analysis; validation; visualization; methodology; writing - original draft; writing - review and editing. **Stefan Prekovic**: investigation; data curation; formal analysis; visualization; writing - review and editing. **Philip Vlummens**: Data curation; writing - review and editing. **Eleni Staessens**: Investigation; writing - review and editing. **Karlien Van Wesemael**: Investigation. **Jonathan Thommis**: Investigation; formal analysis. **Daria Fijalkowska**: data curation; formal analysis; writing - review and editing. **Guillaume Acke**: data curation; formal analysis; visualization; writing - review and editing. **Wilbert Zwart**: writing - review and editing. **Ilse M Beck**: Conceptualization; writing - review and editing; supervision. **Fritz Offner**: writing - review and editing; supervision. **Karolien De Bosscher**: Conceptualization; writing - review and editing; supervision.

## Disclosure and competing interest statement

The authors have no disclosure or competing financial interests.

## Notes

### Competing Interest Statement

The authors have declared no competing interest.

