## Supplemental information for "Crosstalk between the glucocorticoid and mineralocorticoid receptor boosts glucocorticoid-induced killing of multiple myeloma cells"

### Supplementary Information

### Figure S1

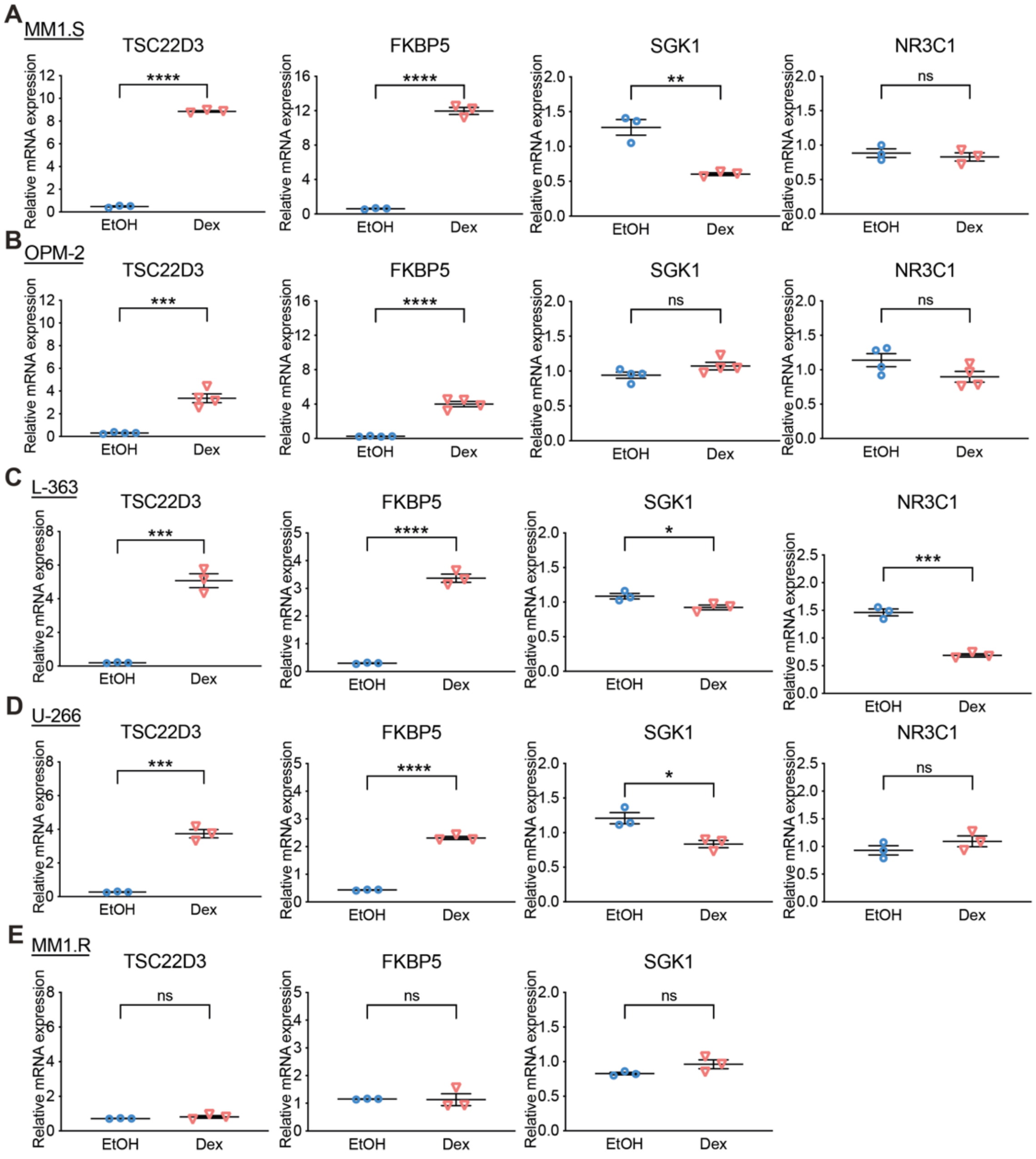


**Fig. S1: Target genes of GR and MR are differentially regulated at the mRNA level between MM cell lines with varying GC responsiveness.**

(**A-E**) Different myeloma cell lines, i.e. (**A**) MM1.S (N=3) (**B**) OPM-2 (N=4), (**C**) L-363 (N=3), (**D**) U-266 (N=3) and (**E**) MM1.R (N=3) cells were treated for 6h with Dex ($10^{-6}$M) or solvent control (EtOH). RNA was isolated and subjected to RT-qPCR analyses, hereby assaying the mRNA levels of *TSC22D3* (GILZ), *FKBP5*, *SGK1* and *NR3C1* (GR). Note that *NR3C1* is not expressed in MM1.R cells. Data analyses were performed using qBaseplus with *SDHA*, *RPL13A* and *YWHAZ* serving as reference genes. The scatter plots represent the mean (solid line) +/- SEM. Statistical analyses were performed using GraphPad Prism 9, using a two-sided unpaired t-test.

### Figure S2

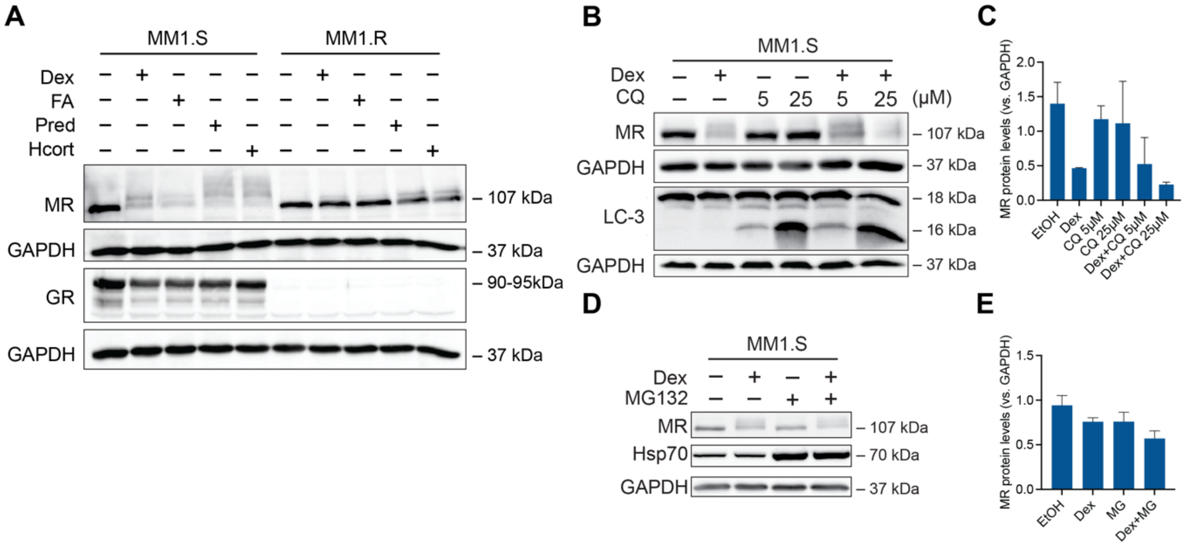


**Fig. S2: MR protein levels are also downregulated by other GCs and neither proteasomal nor lysosomal degradation are involved.**

(**A**) MM1.S and MM1.R cells were treated with Dex, FA, Pred, Hcort ($10^{-6}$M each), and analyzed via WB (N=3).

(**B-C**) MM1.S cells were treated for 24h with solvent, Dex ($10^{-6}$M), CQ ($5.10^{-6}$M or $25.10^{-6}$M), a Dex/CQ combination or solvent control, followed by (**C**) WB analyses (N=2) and (**D**) band densitometric analysis.

(**D-E**) MM1.S cells were treated for 6h with Dex ($10^{-6}$M), MG132 ($10^{-6}$M), a Dex/MG132 combination or solvent control (N=3), followed by (**E**) WB analyses (N=3) and (**F**) band densitometric analysis.

Data information: (**A**, **B, D**) Protein lysates were subjected to WB analysis, determining the protein levels of GR (90-95kDa), MR (107kDa), LC-3 (16-18kDa; positive control for inhibition of lysosomal degradation) or Hsp70 (70kDa; positive control for proteasome inhibition). GAPDH (37kDa) served as a loading control. One representative image for each WB experiment is shown, with the number of biological replicates mentioned in each panel description. (**C, E**) Band densitometric analysis (bar plot) shows the normalized MR protein levels (vs. GAPDH), averaged over the biological experiments (+/- SEM).

### Figure S3

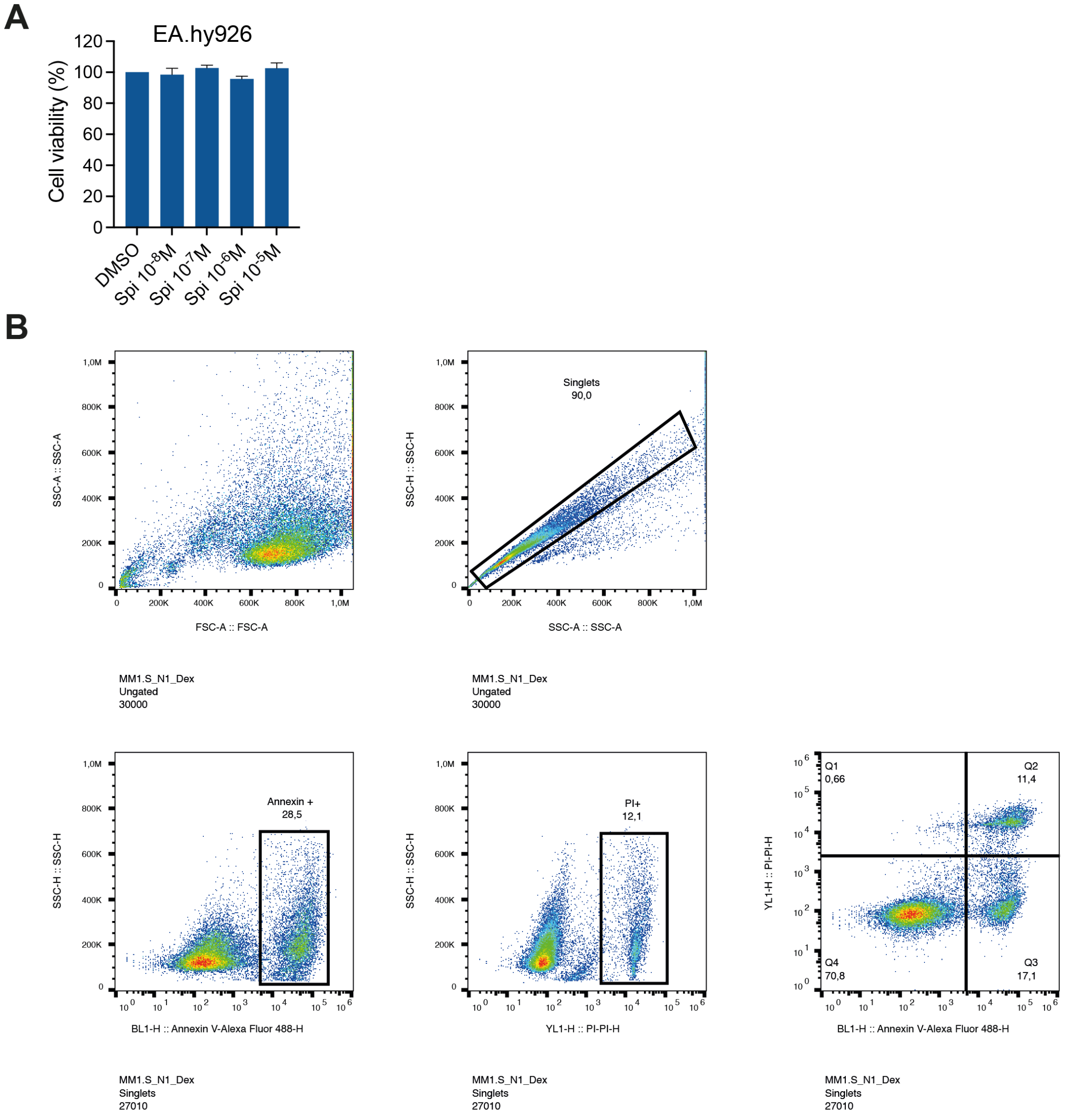


**Fig.S3: EA.hy926 cell-based toxicity control for Spi treatment along with the gating strategy of the flow cytometric analyses in MM1.S cells.**

(**A**) EA.hy926 cells were treated for 72h with a Spi concentration range (10-5M-10-8M), followed by CellTiterGlo cell viability assays (N=3). Solvent control (DMSO) was set at 100% and all other conditions were recalculated accordingly. The bar plot represents the average +/- SEM. Statistical analyses were performed using GraphPad Prism 9, using one-way ANOVA with post-hoc testing, comparing $10^{-8}$M Spi with all other Spi concentrations. No significant differences were found.

(**B**) Gating strategy for Annexin V/PI flow cytometric analyses, illustrated for a Dex-treated sample.

**Figure S4**

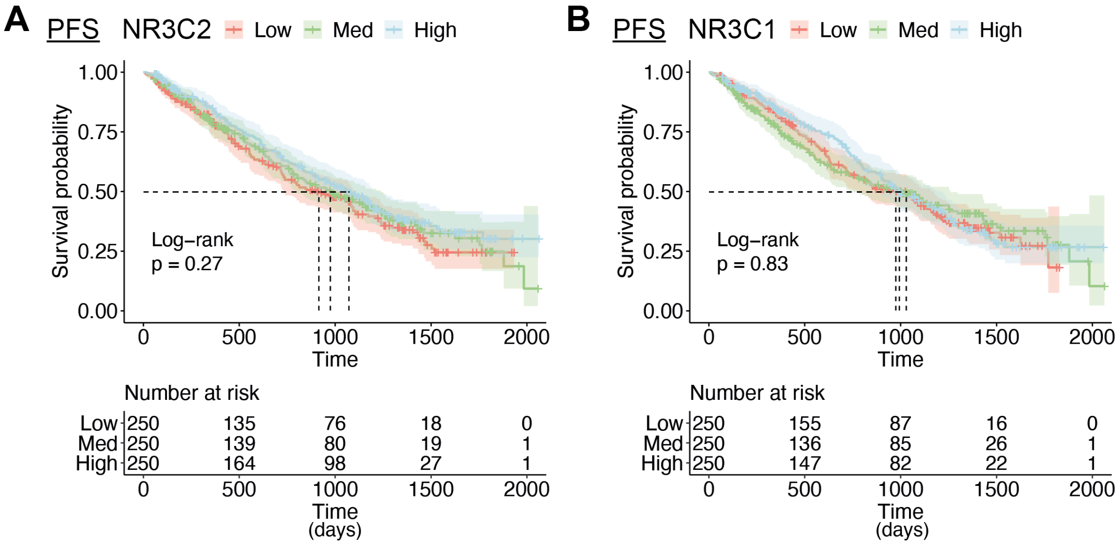


**Fig. S4: Link between NR3C2 or NR3C1 expression levels and survival characteristics.** (**A-B**) Kaplan-Meier curves of the MMRF patient cohort, depicting the survival probability in function of progression-free survival (PFS) for low, medium or high expression of (**A**) *NR3C2* or (**B**) *NR3C1*. Patients were divided in 3 groups based on their expression levels of *NR3C2* or *NR3C1*. Statistical analyses were performed in R (package survival), using a log-rank test.

**Figure S5**

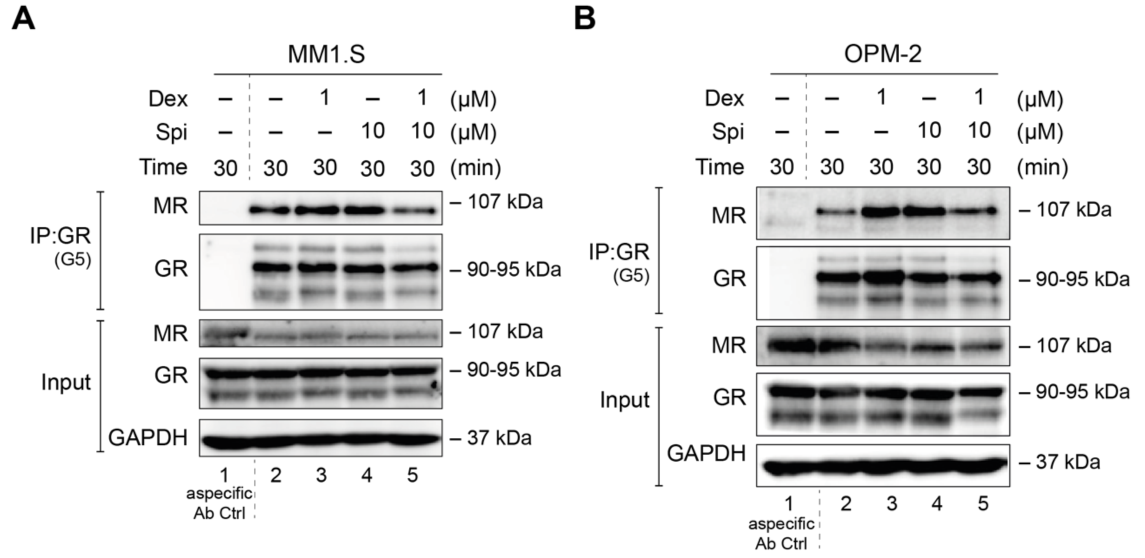


**Fig. S5: Biological replicates of the co-IPs of GR with MR in MM1.S and OPM-2 cell lines.** (**A-B**) Two myeloma cell lines, i.e. (**A**) MM1.S and (**B**) OPM-2 cells, were treated with Dex ($10^{-6}$M), Spi ($10^{-5}$M), a Dex-Spi combination or solvent control for 30min. Protein lysates were prepared and subjected to endogenous IP using GR (G5) antibody. Thereafter, WB analyses were performed to determine co-IP of GR (90-95kDa) with MR (107kDa); GAPDH served as loading control for the input fraction. Lane 1 represents the non-specific antibody control. These WBs represent the second biological replicate of the co-IPs presented in Fig.5D,F.

### Figure S6

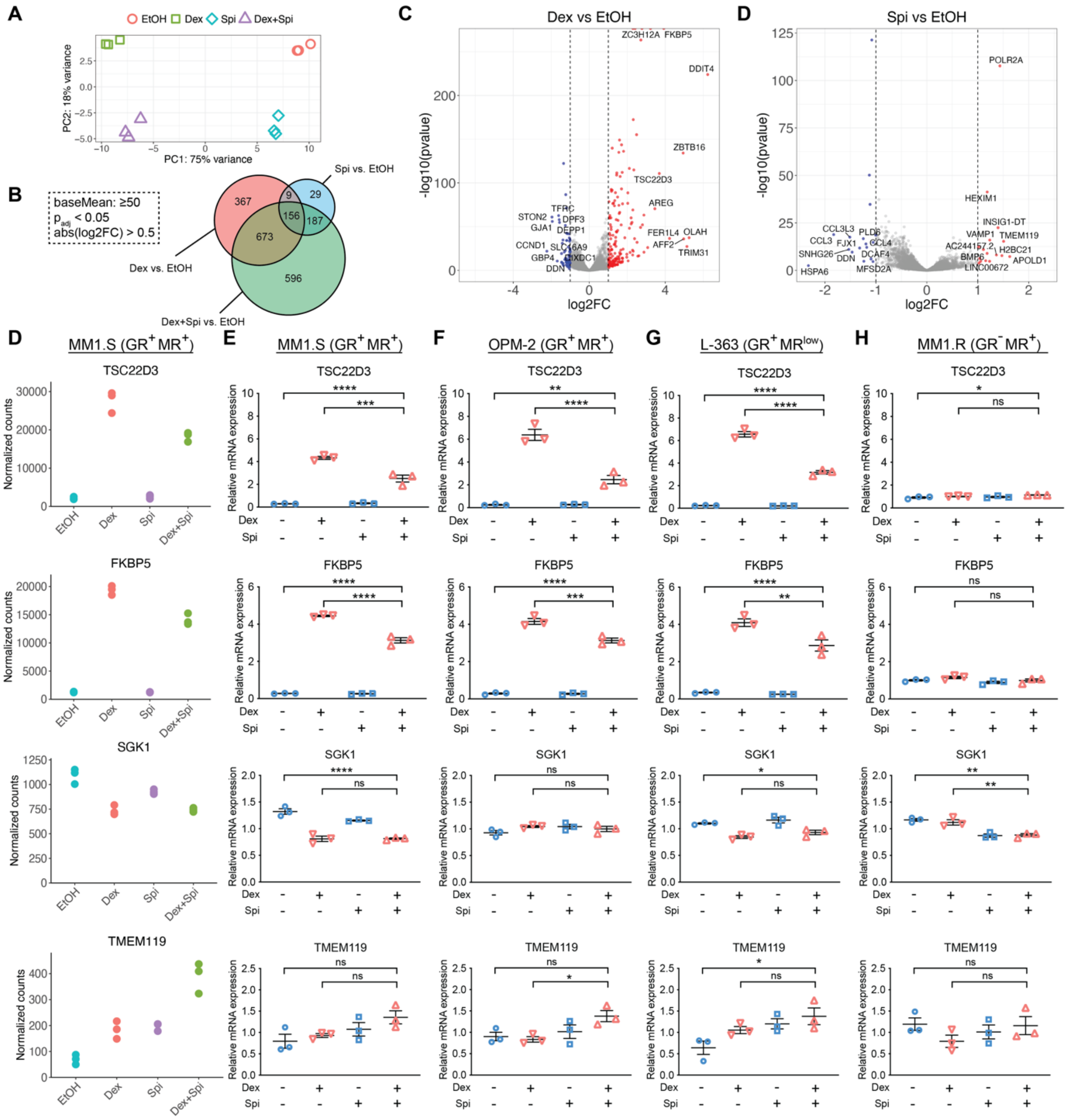

**Fig. S6: Validation of the RNA-sequencing results at the mRNA level in several myeloma cell lines.**

(**A-D**) MM1.S cells were treated with Dex ($10^{-6}$M), Spi ($10^{-5}$M), a Dex/Spi combination or solvent control (EtOH) for 6h, followed by RNA-seq analysis. (**A**) Principal component analysis of each condition and biological replicate of the RNA-seq experimental setup (6h treatment, N=3).

(**B**) Venn-diagram depicting the number of protein coding genes with baseMean≥50 that are significantly regulated ($p_{adj}$<0.05) and have an abs(log2FC)>0.5 in and between different pairwise comparisons (Dex-Spi vs EtOH; Dex vs EtOH; Spi vs EtOH).

(**C, D**) Volcano plots depicting the $p_{adj}$ (log10 scale) in function of the log2FC for all genes with baseMean ≥50 for the pairwise comparison (**C**) Dex vs EtOH or (**D**) Spi vs EtOH. Significantly regulated genes ($p_{adj}$<0.05) are colored in red when log2FC>1 or blue when log2FC<-1; non-significant genes ($p_{adj}$>0.05) in grey. The gene names are displayed for those genes having the largest abs(log2FC) values (top 10 upregulated/downregulated).

(**D**) The normalized counts are plotted for several genes identified by RNA-sequencing in MM1.S.

(**E-H**) Several myeloma cell lines, i.e. (**E**) MM1.S, (**F**) OPM-2, (**G**) L-363 and (**H**) MM1.R cells were treated with Dex ($10^{-6}$M), Spi ($10^{-5}$M), a Dex/Spi combination or solvent control (EtOH) for 6h (all cell lines N=3). RNA isolation and RT-qPCR analyses were performed to determine the mRNA expression levels of *TSC22D3* (GILZ), *FKBP5*, *SGK1* and *TMEM119*. Data analyses were performed using qBaseplus with *SDHA*, *RPL13A* and *YWHAZ* serving as reference genes. The scatter plots represent the mean +/- SEM (N=3 for all cell lines). Statistical analyses were performed using GraphPad Prism 9, using a one-way ANOVA with post-hoc testing.

### Figure S7

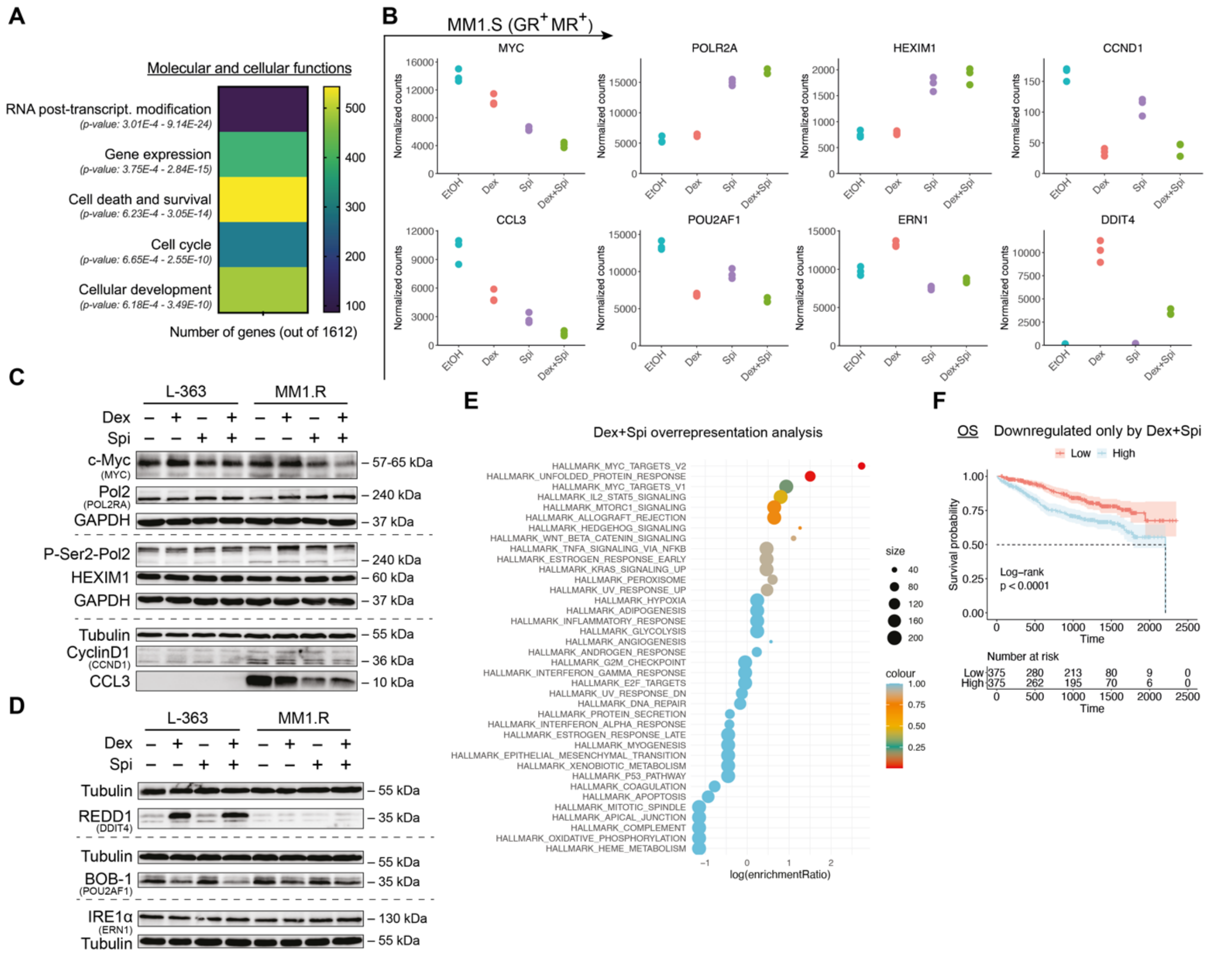

**Fig. S7: Validation of the RNA-sequencing results at the protein level in several myeloma cell lines along with functional annotations.**

(**A**) Molecular and cellular functions identified via core analyses in IPA using protein coding genes with baseMean≥50, $p_{adj}$<0.05 and abs(log2FC)≥0.5 as input for the pairwise comparisons Dex-Spi vs EtOH. The number of genes that are allocated to a certain molecular and cellular function are depicted as a heatmap and the corresponding p-value range is mentioned.

(**B**) The normalized counts are plotted for several genes identified by RNA-sequencing in MM1.S.

(**C**) L-363 and MM1.R cells were treated with Dex ($10^{-6}$M), Spi ($10^{-5}$M), a Dex/Spi combination or solvent control (EtOH) for 24h (both N=3). Protein lysates were prepared and subjected to WB analyses, hereby assessing the protein levels of c-myc (MYC, 57-65kDa), (P-Ser2) RNA-Pol2 (POLR2A, 240kDa), HEXIM (HEXIM1, 60kDa), cyclinD1 (CCND1, 36kDa) and MIP-1α (CCL3, 10kDa).

(**D**) L-363 and MM1.R cells were treated with Dex ($10^{-6}$M), Spi ($10^{-5}$M), a Dex/Spi combination or solvent control (EtOH) for 24h (both N=3). Protein lysates were prepared and subjected to WB analyses, hereby assessing the protein levels of REDD1(DDIT4, 35kDa), BOB-1 (POU2AF1, 35kDa) and IRE1α (ERN1, 110-130kDa). Tubulin (55kDa) served as a loading control.

(**E**) GSEA-based overrepresentation analysis for the genes uniquely downregulated by Dex-Spi, hereby identifying hallmarks that are enriched. Color refers to the significance level (p-value) and size to the number of enriched genes per hallmark.
(**F**) Kaplan-Meier curve of the CoMMpass patient cohort (N=750), depicting the survival probability in function of overall survival (OS) for low or high expression of genes that were uniquely downregulated by the Dex-Spi combination. Statistical analyses were performed in R (package survival), using a log-rank test.

Data information: (**B, D**) One representative image for each WB experiment is shown, with the number of biological replicates mentioned in each panel description.

**Figure S8**

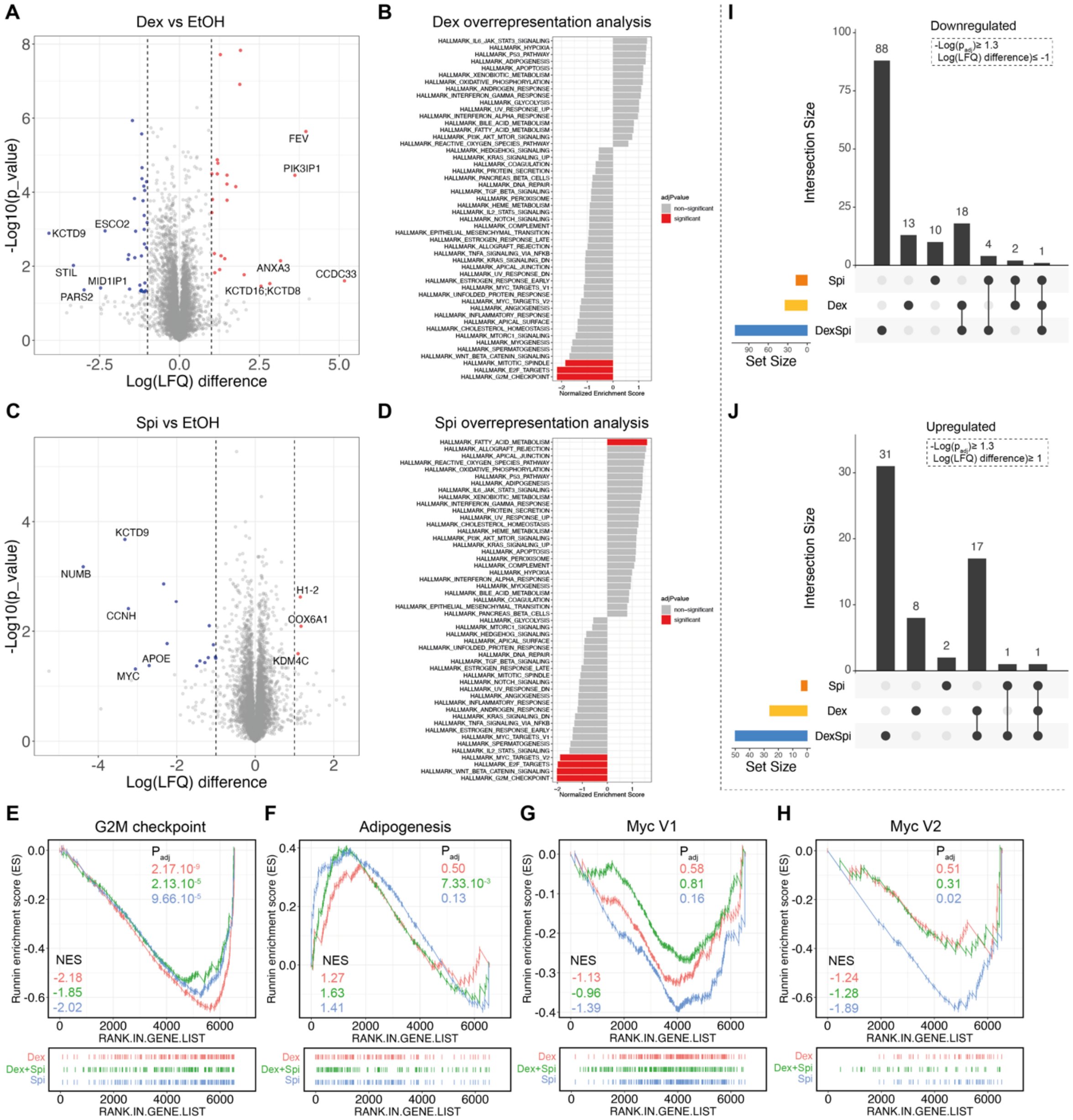

**Fig. S8: MS-based shotgun proteomics of other pairwise comparisons along with additional single hallmark plots enriched upon Dex-Spi treatment.**

(**A, C**) Volcano plots depicting the $p_{adj}$ (log10 scale) in function of the log(LFQ) in the pairwise comparison (**A**) Dex vs EtOH and (**C**) Spi vs EtOH. Significantly regulated proteins -log(padj)≥1.3 are colored in red (log(LFQ)>1, upregulated) or blue (log(LFQ)<-1, downregulated); non-significant genes (-log($p_{adj}$)<1.3) in grey.

(**B, D**) GSEA-based overrepresentation analysis for the proteins regulated by (**B**) Dex or (**D**) Spi, hereby identifying hallmarks that are significantly (red) or non-significantly (grey) enriched.

(**E-H**) GSEA of single hallmarks, i.e. (**E**) G2M checkpoint (**F**) Adipogenesis, (**G**) Myc V1 (**H**) Myc V2 targets, for each pairwise comparison, along with the respective normalized enrichment score (NES) and $p_{adj}$.

(**I-J**) Upset plots depicting the number of proteins that are (**I**) downregulated or (**J**) upregulated in and between pairwise comparisons (Dex-Spi vs EtOH, Dex vs EtOH and Spi vs EtOH).

**Figure S9**

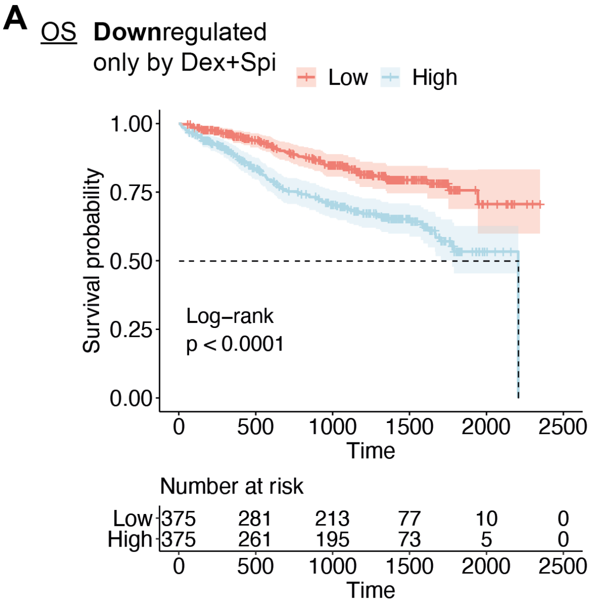


**Fig. S9: OS analysis of the CoMMpass cohort using differentially regulated proteins as input. (A-B)** Kaplan-Meier curve of the CoMMpass patient cohort (N=750), depicting the survival probability in function of overall survival (OS) for low or high expression of proteins that were uniquely downregulated by the Dex-Spi combination. Statistical analyses were performed in R (package survival), using a log-rank test.

**Table S1. RNA-seq-derived interaction term genes.** The baseMean, log2FC and $p_{adj}$ of these genes are mentioned.

| Gene | baseMean | Log2FC | $p_{adj}$ |
|---|---|---|---|
| STK38L | 4064.5 | 0.57 | 1.46E-09 |
| TFRC | 2334.4 | 0.46 | 4.80E-02 |
| CCR1 | 3376.0 | 0.38 | 3.10E-03 |
| SCD | 13024.2 | 0.38 | 4.80E-02 |
| BARX2 | 2207.5 | 0.34 | 1.86E-02 |
| JCHAIN | 83330.9 | 0.34 | 1.25E-02 |
| CD28 | 6686.1 | 0.32 | 2.80E-02 |
| POU2AF1 | 9033.7 | 0.31 | 3.51E-02 |
| PLXNC1 | 7038.2 | 0.31 | 1.46E-03 |
| METTL7A | 16737.2 | 0.25 | 4.69E-02 |
| PM20D2 | 3878.5 | 0.23 | 2.85E-02 |
| PFKFB2 | 5155.7 | -0.25 | 4.86E-02 |
| ERN1 | 9782.2 | -0.27 | 3.45E-02 |
| GLUL | 5023.7 | -0.31 | 2.85E-02 |
| TREML2 | 3849.4 | -0.31 | 8.37E-03 |
| SLFN5 | 1586.6 | -0.31 | 3.90E-02 |
| SMC4 | 22296.2 | -0.32 | 4.54E-03 |
| BACH2 | 1608.1 | -0.35 | 3.22E-02 |
| CELF2 | 12208.6 | -0.36 | 1.04E-02 |
| CYSLTR1 | 2308.5 | -0.37 | 2.44E-02 |
| PLEKHA7 | 3633.0 | -0.37 | 7.16E-03 |
| IRAG2 | 2114.1 | -0.38 | 1.16E-02 |
| FKBP5 | 9001.7 | -0.42 | 1.46E-03 |
| GCSAM | 2554.6 | -0.44 | 8.31E-03 |
| SESN1 | 3963.5 | -0.44 | 3.10E-03 |
| SLC38A2 | 6626.6 | -0.45 | 4.54E-03 |
| MEI1 | 2981.3 | -0.46 | 9.73E-03 |
| INSR | 2944.2 | -0.46 | 5.56E-05 |
| HMGB3 | 2430.2 | -0.50 | 2.23E-06 |
| TMSB4X | 16052.7 | -0.52 | 3.66E-05 |
| ZC3H12A | 3422.8 | -0.73 | 1.08E-08 |
| NUDT16 | 2981.9 | -0.79 | 8.62E-17 |
| RHOB | 1914.3 | -0.82 | 1.20E-04 |
| TSC22D3 | 12654.5 | -0.82 | 3.45E-02 |
| TXNIP | 19898.5 | -1.02 | 1.08E-09 |
| FBXO32 | 2535.1 | -1.10 | 5.12E-14 |
| DDIT4 | 3500.6 | -1.67 | 5.24E-07 |

**Table S2. siRNA target sequence identifiers.**

| Gene (protein) | siRNA information |
|---|---|
| NR3C1 (GR) | M-003424-03-0010, siGENOME Human NR3C1 siRNA – SMARTpool (mixture of 4 siRNA's):<br>• siGENOME SMARTpool siRNA D-003424-04 NR3C1<br>• siGENOME SMARTpool siRNA D-003424-06 NR3C1<br>• siGENOME SMARTpool siRNA D-003424-19 NR3C1<br>• siGENOME SMARTpool siRNA D-003424-20 NR3C1 |
| NR3C2 (MR) | M-003425-02-0010, siGENOME Human NR3C2 siRNA – SMARTpool (mixture of 4 siRNA's):<br>• siGENOME SMARTpool siRNA D-003425-01 NR3C2<br>• siGENOME SMARTpool siRNA D-003425-02 NR3C2<br>• siGENOME SMARTpool siRNA D-003425-04 NR3C2<br>• siGENOME SMARTpool siRNA D-003425-05 NR3C2 |
| Non-targeting (Control) | D-001206-13-20, siGENOME Non-targeting siRNA Pool 1 |

**Table S3. Sequences of qPCR primers.**

| Target | Primer forward | Primer reverse |
|---|---|---|
| MR | CAGGGGATGCACCAAATCAG | AGGCCATCCTTTGGAATTGTG |
| GR | TGATGAAGCTTCAGGATGTCA | TTCGAGCTTCCAGGTTCATTC |
| FKBP5 | AGTAGAAATCCACCTGGAAGGC | ATTTAGGCTTCCCTGCCTCT |
| TSC22D3 | GCGTGAGAACACCCTGTTGA | TCAGACAGGACTGGAACTTCTCC |
| SGK1 | GAGATTGTGTTAGCTCCAAAGC | CTGTGATCAGGCATACCACACT |
| TMEM119 | GACCCCTGCACACATTACGA | TGTTTCCGTAGAGTGCCTCG |

**Table S4. Primary antibodies.**

| Target | Catalog number | Obtained from |
|---|---|---|
| MR 6G1 | NA | Dr. Gomez-Sanchez (Univ. of Mississippi) |
| GR H300 | sc-8992 | Santa Cruz Biotechnology |
| GR G5 | sc-393232 | Santa Cruz Biotechnology |
| PARP | 556494 | BD Biosciences |
| Cleaved-caspase 3 | 9664 | Cell Signaling |
| Bim | sc-374358 | Santa Cruz Biotechnology |
| Bcl-XL | sc-8392 | Santa Cruz Biotechnology |
| β-catenin | C7202 | Sigma |
| LC-3 | L8918 | Sigma |
| Hsp70 | ADI-SPA-810 | Enzo life sciences |
| RNA pol2 | sc899 | Santa Cruz Biotechnology |
| RNA pol2 (P-Ser2) | ab5095 | Abcam |
| c-myc | CST5605T | Cell signaling |
| HEXIM | CST12604S | Cell signaling |
| Cyclin-D1 | CST5506S | Cell signaling |
| CCL3 | ab259372 | Abcam |
| REDD1 | 10638-1-AP | Protein tech |
| IRE1α | sc-390960 | Santa Cruz Biotechnology |
| BOB-1 | CST43079 | Cell signaling |
| GAPDH | ab9485 | Abcam |
| GAPDH | G8795 | Sigma |
| Tubulin | T5168 | Sigma |
